# Characterization of the apicomplexan amino acid transporter (ApiAT) family in *Toxoplasma gondii*

**DOI:** 10.1101/306993

**Authors:** Kathryn E. R. Parker, Stephen J. Fairweather, Esther Rajendran, Martin Blume, Malcolm J. McConville, Stefan Bröer, Kiaran Kirk, Giel G. van Dooren

## Abstract

Apicomplexan parasites are auxotrophic for a range of amino acids which must be salvaged from their host cells, either through direct uptake or degradation of host proteins. Here, we describe a family of plasma membrane-localized amino acid transporters, termed the Apicomplexan Amino acid Transporters (ApiATs), that are ubiquitous in apicomplexan parasites. Functional characterization of the ApiATs of *Toxoplasma gondii* indicate that several of these transporters are important for intracellular growth of the tachyzoite stage of the parasite, which is responsible for acute infections. We demonstrate that the ApiAT protein *Tg*ApiAT5-3 is an exchanger for aromatic and large neutral amino acids, with particular importance for L-tyrosine scavenging and amino acid homeostasis, and that *Tg*ApiAT5-3 is critical for parasite virulence. Our data indicate that *T. gondii* expresses additional proteins involved in the uptake of aromatic amino acids, and we present a model for the uptake and homeostasis of these amino acids. Our findings identify a family of amino acid transporters in apicomplexans, and highlight the importance of amino acid scavenging for the biology of this important phylum of intracellular parasites.

**Author Summary:** The Apicomplexa comprise a large number of parasitic protozoa that have obligate intracellular lifestyles and cause significant human and animal diseases, including malaria, cryptosporidiosis, toxoplasmosis, coccidiosis in poultry, and various cattle fevers. Apicomplexans must scavenge essential nutrients from their hosts in order to proliferate and cause disease, including a range of amino acids. The direct uptake of these nutrients is presumed to be mediated by transporter proteins located in the plasma membrane of intracellular stages, although the identities of these proteins are poorly defined. Using a combination of bioinformatic, genetic, cell biological, and physiological approaches, we have characterized a family of plasma membrane-localized transporter proteins that we have called the Apicomplexan Amino acid Transporters (ApiATs). The family is found in apicomplexans and their closest free-living relatives. We show that *Tg*ApiAT5-3, a member of the family in the apicomplexan *Toxoplasma gondii*, is an exchanger for aromatic and large neutral amino acids. In particular, it is critical for uptake of tyrosine, and for parasite virulence in a mouse infection model. We conclude that ApiATs are a family of plasma membrane transporters that play crucial roles in amino acid scavenging by apicomplexan parasites.

## Introduction

Apicomplexans are intracellular parasites that cause a range of diseases in humans and animals, imparting a major health and economic burden on many countries. In humans, *Plasmodium* species are the causative agents of malaria, while *Cryptosporidium* is a major cause of diarrheal disease and death in children in the developing world (1). *Toxoplasma gondii* can infect virtually all nucleated cells in warm-blooded animals, and is thought to chronically infect one-third of the world’s human population. *T. gondii* infections are usually asymptomatic, but infection in immunocompromised patients may lead to life-threatening toxoplasmic encephalitis, and congenital toxoplasmosis may result in severe birth defects or death of the developing fetus (2).

A common feature of parasites is that they rely on their hosts to supply them with the nutrients necessary for their growth and replication, such as sugars, amino acids, nucleosides, and vitamins. Transporters are integral membrane proteins that facilitate the transfer of substrates across biological membranes. In apicomplexans, transporters provide the major route for the acquisition of nutrients and the removal of waste products across the plasma membrane (3), and these proteins are important for parasite survival and virulence (4, 5). Despite this, the transporters responsible for the uptake of many essential nutrients in apicomplexans have not been defined.

A family of Novel Putative Transporters (the NPT family) was initially identified in *Plasmodium falciparum* using a bioinformatics approach (6). The five *P. falciparum* NPT family proteins were predicted to be polytopic membrane proteins with a secondary structure characteristic of solute transporters, although they have limited sequence similarity to other eukaryotic or prokaryotic transporters. The NPT family protein *Pb*NPT1 localizes to the plasma membrane of the mouse malaria-causing parasite *P. berghei*, and is essential for gametocyte development in the murine host and subsequent mosquito transmission (4, 5, 7). Other *P. berghei* NPT family proteins, *Pb*MFR4 and *Pb*MFR5, are essential for progression through the insect stages of the life cycle, while *Pb*MFR2 and *Pb*MFR3 are important for exflagellation of male gametes and sporozoite formation, respectively, but are not essential for completion of the *P. berghei* life cycle (4). In *T. gondii, Tg*NPT1 localizes to the plasma membrane and is essential for parasite growth and virulence (5). Both *Pb*NPT1 and *Tg*NPT1 are cationic amino acid transporters, with *Pb*NPT1 functioning as a general cationic amino acid transporter and *Tg*NPT1 functioning as a selective arginine transporter (5). The functions of other NPT-family proteins are not known, although one member of the family has been associated with susceptibility to the anti-*T. gondii* drug sinefungin (8).

In this study, we have demonstrated that the NPTs are phylogenetically related, and broadly distributed within the apicomplexan phylum and their closest free-living relatives. We have characterized the NPT family proteins in *T. gondii*, demonstrating that 10 of the sixteen members of the family are expressed in the disease-causing tachyzoite stage of the parasite, and that the majority of these localize to the parasite plasma membrane. We have demonstrated that at least three of these proteins are important for *in vitro* growth of the parasite. Using a combination of genetic, physiological and heterologous expression approaches, we have shown that one of the previously uncharacterized *T. gondii* NPT-family members transports aromatic and large neutral amino acids, and that this transporter is particularly important for the uptake of tyrosine into the parasite. We conclude that NPTs are a family of amino acid transporter proteins found in apicomplexans, and we propose that the family be renamed the *<u>Api</u>*complexan *<u>A</u>*mino acid *<u>T</u>*ransporter (ApiAT) family.

## Results

### ApiATs are broadly-distributed in apicomplexan parasites

To identify ApiAT-family proteins in the apicomplexan parasites *T. gondii, Neospora caninum, Eimeria tenella, P. falciparum, P. berghei, Theileria annulata, Babesia bovis* and *Cryptosporidium parvum*, we undertook Basic Local Alignment Search Tool (BLAST) searches using *Pb*NPT1 as an initial query sequence (www.eupathdb.org; (9)) We also undertook BLAST searches of the genomes from the chromerids *Chromera velia* and *Vitrella brassicaformis*, which are close free-living relatives of apicomplexans (10), and a broad range of other eukaryotes (www.eupathdb.org, www.blast.ncbi.nlm.nih.gov; (9, 11)). In addition to the previously described five *Plasmodium* ApiAT family proteins, we identified sixteen ApiAT family proteins in both *T. gondii* and *N. caninum*, nine in *E. tenella*, six in *T. annulata*, five in *B. bovis*, three in *V. brassicaformis*, and one each in *C. parvum* and *C. velia* (Fig S1). Using this strategy, we were unable to identify ApiAT family proteins outside of the apicomplexan/chromerid lineage. Using *Pb*NPT1 as a search query in profile hidden Markov Model searches, we identified the LAT3 and LAT4 proteins from humans (http://hmmer.org; (12); Fig S2). LAT3 and LAT4 are members of the SLC43 family of the major facilitator superfamily of transporters, and mediate the transport of branched chain and other large neutral amino acids (13).

To determine the relationships between ApiAT family proteins, we constructed a multiple sequence alignment. This revealed the presence of a major facilitator superfamily (MFS) signature sequence between transmembrane domains 2 and 3 of most ApiAT family protein (Fig S1; (14, 15)). Most ApiAT proteins were predicted to be polytopic membrane proteins containing 12 transmembrane domains (www.cbs.dtu.dk/services/TMHMM/; (16)), and exhibited highest sequence similarity in the regions encompassing these transmembrane domains (Fig S1). These analyses are consistent with previous studies and protein database annotations placing members of the ApiAT family into the major facilitator superfamily of transporters (5, 6).

We next performed a maximum likelihood phylogenetic analysis. This revealed the presence of multiple ApiAT subfamilies (Fig 1). Orthologs of the ApiAT2 subfamily were present in all organisms in the study, with the exception of *Cryptosporidium parvum* and the free living Chromerid species (Fig 1). Members of the ApiAT3, ApiAT5, ApiAT6 and ApiAT7 subfamilies were restricted to coccidians (a group of apicomplexans that includes *T. gondii* and *N. caninum*), and the ApiAT9 family was restricted to the piroplasms *T. annulata* and *B. bovis* (Fig 1). *Plasmodium* ApiAT3 branched with the coccidian ApiAT3 subfamily, although bootstrap support for this association was weak (Fig 1). The ApiAT8 subfamily comprises two members in each *Plasmodium* species examined. This family includes the previously described cationic transporter *Pb*NPT1 (here annotated as *Pb*ApiAT8-1). Although similar in function to the *T. gondii* arginine transporter *Tg*ApiAT1 (previously *Tg*NPT1), *Pb*ApiAT8-1 and *Tg*ApiAT1 appear not to be orthologous.

**Fig 1.**
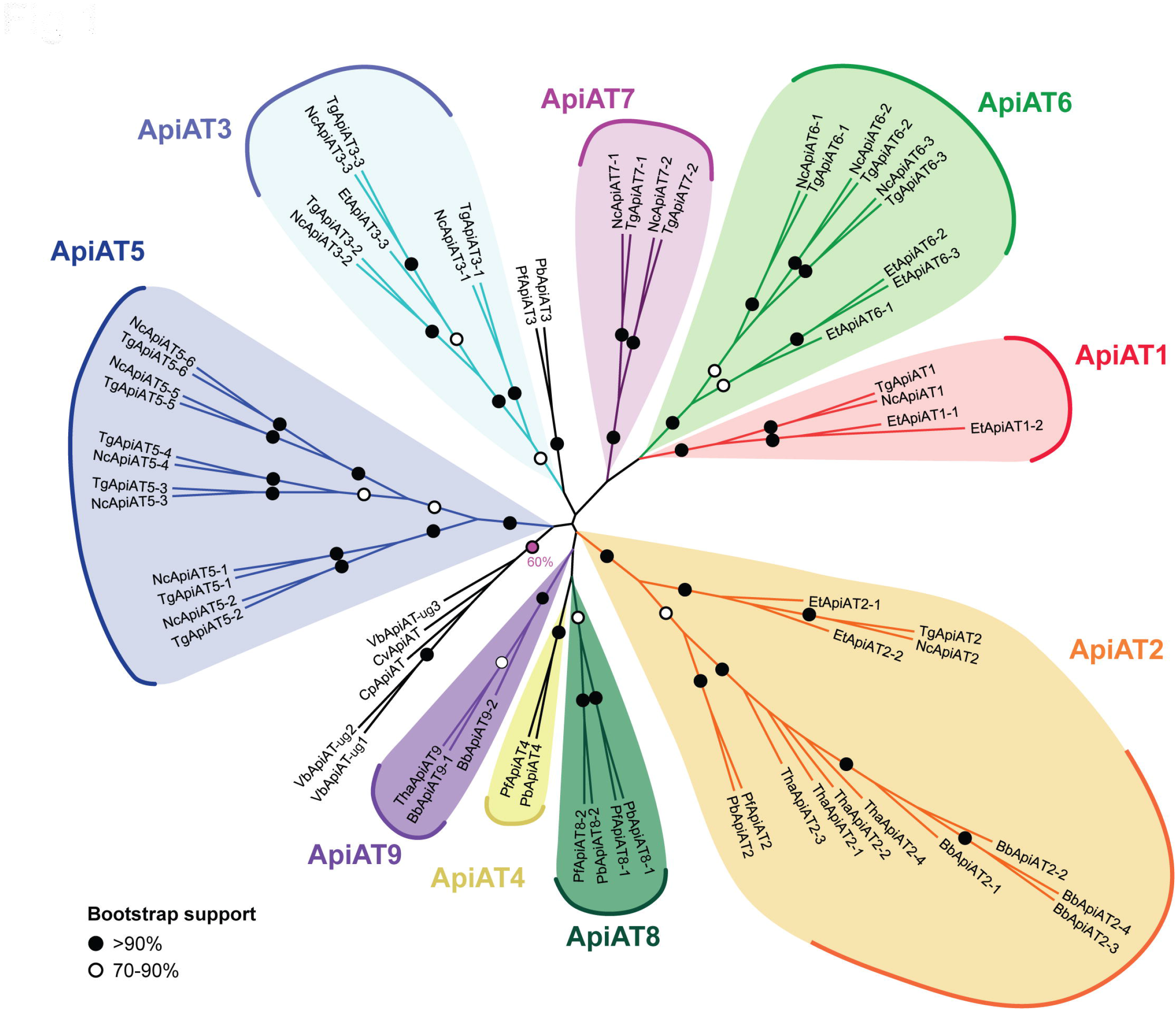
Phylogenetic analysis of ApiAT family proteins. Consensus maximum likelihood tree of ApiAT family proteins. The tree was generated from a multiple sequence alignment of 67 putative ApiAT proteins from a range of apicomplexans and chromerids, with 452 residues used in the analysis. Bootstrap values are depicted by black circles (>90% support), white circles (70-90% support), or a pink circle (60% for a group consisting of *Cryptosporidium* and chromerid proteins). The tree is unrooted. Abbreviations: Bb, *Babesia bovis*; Cp, *Cryptosporidium parvum*; Cv, *Chromera velia*; Et, *Eimeria tenella*; Nc, *Neospora caninum*; Pb. *Plasmodium berghei*; Pf, *Plasmodium falciparum*; Tg, *Toxoplasma gondii*; Tha, *Theileria annulata*; Vb, *Vitrella brassicaformis*.

### *T. gondii* ApiATs localize to the parasite periphery

Previous studies demonstrated that the *P. berghei* ApiAT8-1 protein (previously *Pb*NPT1) localized to the periphery of the parasite (likely to the plasma membrane), and that the *T. gondii* ApiAT1 protein (previously *Tg*NPT1) localized to the plasma membrane (5, 7). To determine the expression pattern and localization of ApiAT family proteins in *T. gondii*, we introduced a hemagglutinin (HA) tag into the 3 ’ end of the open reading frame of the remaining fifteen ApiAT genes (Fig S3A-B).

Western blotting indicated that *Tg*ApiAT2, *Tg*ApiAT3-1, *Tg*ApiAT3-2, *Tg*ApiAT3-3, *Tg*ApiAT5-3, *Tg*ApiAT6-1, *Tg*ApiAT6-2, *Tg*ApiAT6-3, and *Tg*ApiAT7-2 proteins were expressed in tachyzoite stage parasites (Fig 2A-E). We were unable to detect expression of *Tg*ApiAT5-1, *Tg*ApiAT5-2, *Tg*ApiAT5-4, *Tg*ApiAT5-5, *Tg*ApiAT5-6 and *Tg*ApiAT7-1. Immunofluorescence assays (IFAs) demonstrated that *Tg*ApiAT2, *Tg*ApiAT3-1, *Tg*ApiAT3-2, *Tg*ApiAT3-3, *Tg*ApiAT5-3, *Tg*ApiAT6-1 (as reported previously; (17)), and *Tg*ApiAT6-3 localized to parasite periphery, overlapping with the plasma membrane marker P30 (Fig 2F-I). *Tg*ApiAT3-3 showed additional localization to the trans-Golgi network (Fig 2G).

**Fig 2.**
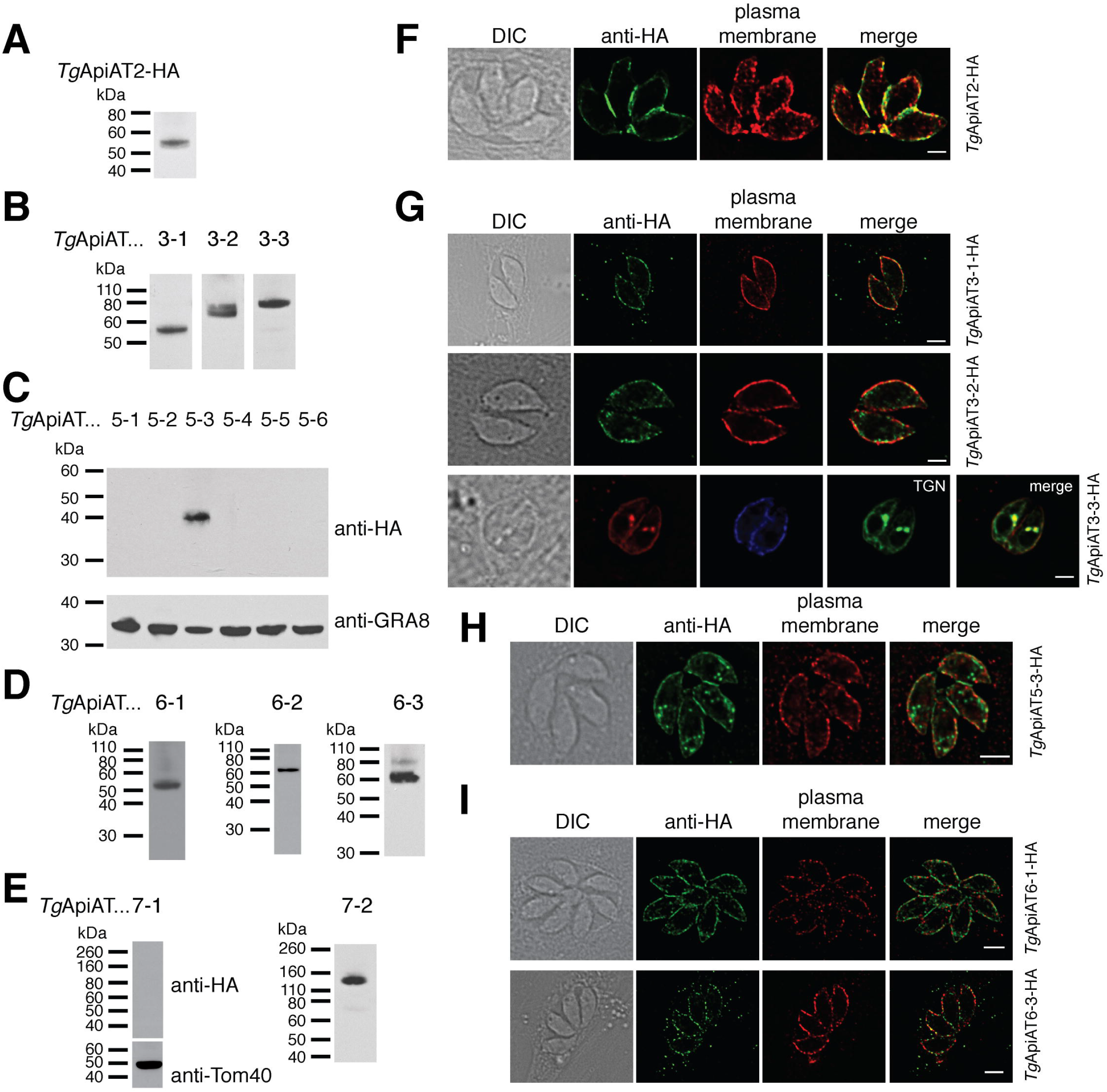
Expression and localization analysis of *T. gondii* ApiAT family proteins. (A-E) Western blots with anti-HA antibodies to measure the expression and molecular mass of tagged *Tg*ApiAT proteins in tachyzoites stages of the parasite. Western blots with antibodies against GRA8 and Tom40 were used to test for the presence of protein in samples where the HA-tagged *Tg*ApiAT protein was not detected. (F-I) Immunofluorescence assays with anti-HA antibodies to determine the localisation of *Tg*ApiAT proteins (green). Samples were co-labelled with antibodies against the plasma membrane marker P30 (red). *Tg*ApiAT3-3-HA-expressing parasites were co-transfected with the trans-Golgi network (TGN) marker Stx6-GFP (56), and labelled with anti-HA (red), anti-P30 (blue) and anti-GFP (green) antibodies. All scale bars are 2 μm.

Although detectable by western blotting (Fig 2D-E), we could not detect *Tg*ApiAT6-3 or *Tg*ApiAT7-2 by IFA, possibly because the level of expression of these proteins was below the detection limits of IFAs. We conclude that ten of the sixteen *Tg*ApiAT proteins are expressed in the tachyzoite stage of *T. gondii*, and those with detectable expression by IFA all localize to the plasma membrane of the parasite.

To determine the importance of *Tg*ApiATs for parasite growth, we attempted to genetically disrupt all sixteen *T. gondii* ApiATs using a CRISPR/Cas9-based approach (18). Using this strategy, we were able to disrupt fifteen of the sixteen *Tg*ApiAT genes (Table S1). *Tg*ApiAT1 could only be disrupted when parasites were grown in excess arginine, as described previously (5). We were unable to generate frameshift mutations in *Tg*ApiAT6-1 after screening 12 clones from three separate transfections of a guide RNA targeting the *Tg*ApiAT6-1 locus. Of these clones, two had a 3 bp insertion and one had a 3 bp deletion, indicating that the guide RNA was capable of targeting the *Tg*ApiAT6-1 locus (Table S1).

To determine which *Tg*ApiATs were important for parasite growth, we performed plaque assays on each of the *Tg*ApiAT knockout (Δ*apiAT*) lines grown in human foreskin fibroblasts (HFFs) and cultured in Dulbecco’s modified Eagle’s medium (DMEM). Compared to parental wild type (WT) controls, we observed greatly reduced plaque sizes in the Δ*apiAT2* and Δ*apiAT5-3*strains (Fig 3). As described previously (5), no plaques were observed in the Δ*apiAT1* strain grown in DMEM (containing 400 μM L-arginine) but normal growth of this strain was observed when grown in Roswell Park Memorial Institute 1640 (RPMI) medium (containing 1.15 mM L-Arg; Fig 3A). By contrast, the remaining 12 Δ*apiAT* lines exhibited plaques that were similar in size to WT controls (Fig 3). To test whether the growth defect in the Δ*api*AT2 strain was due specifically to disruption of the *Tg*ApiAT2 locus, we complemented Δ*api*AT2 with constitutively expressed *Tg*ApiAT2. This restored parasite growth (Fig 3B). These results indicate that *Tg*ApiAT1, *Tg*ApiAT2 and *Tg*ApiAT5-3 are required for normal intracellular growth of *T. gondii* in standard *in vitro* culture conditions.

**Fig 3.**
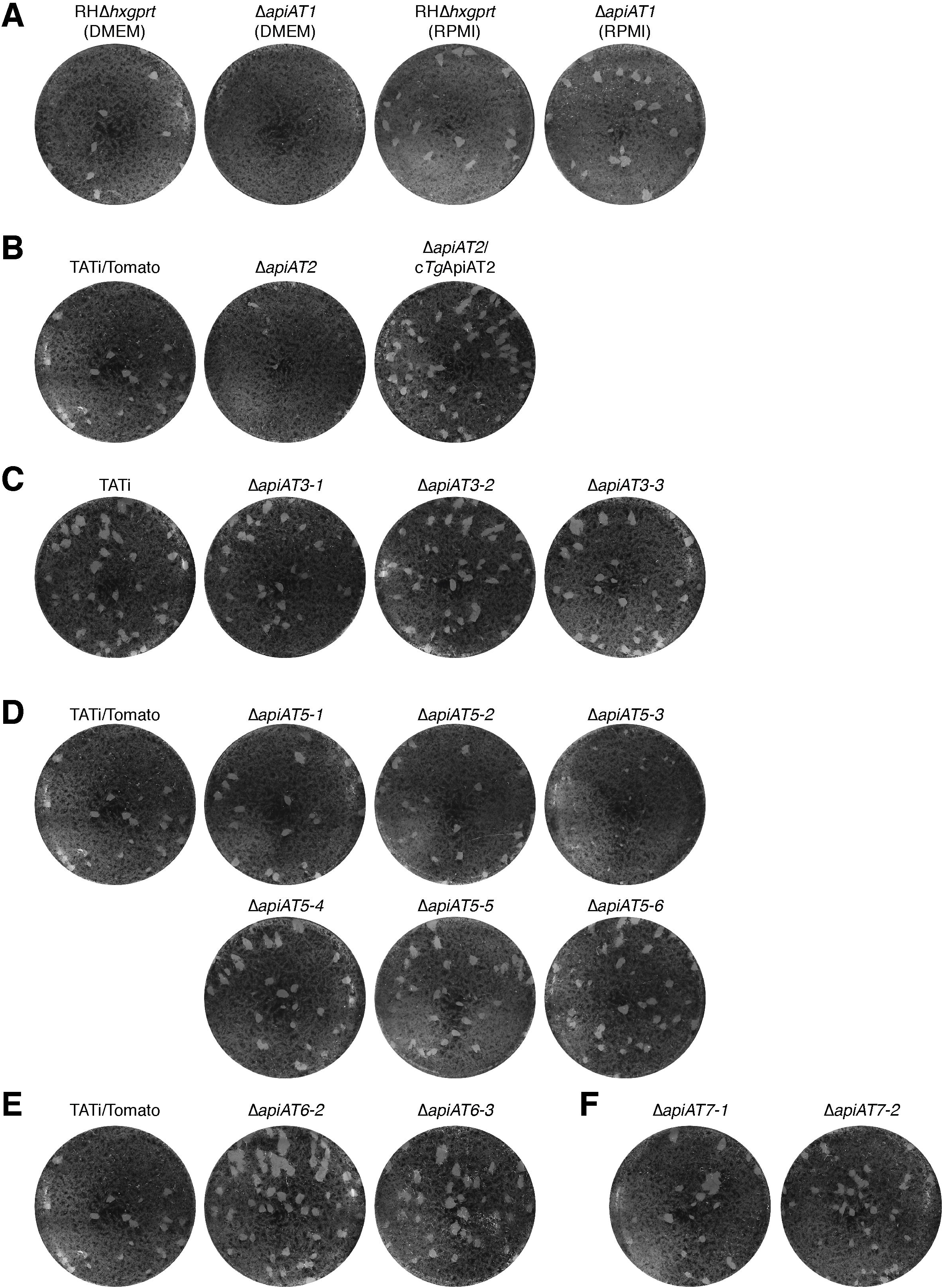
Genetic disruption of *T. gondii* ApiAT family proteins reveals the importance of *Tg*ApiAT2 and *Tg*ApiAT5-3 for parasite growth *in vitro*. (A-F) Plaque assays depicting growth of *Tg*ApiAT knockout strains and their corresponding parental WT strain. 150 parasites were added to wells of a 6-well plate and cultured for 9 days in DMEM g5ies(unless otherwise indicated). (A) WT (RHΔ*hxpgrt*) and *ΔapiAT1* parasites grown in DMEM (left) or RPMI (right). (B) WT (TATi/Tomato), Δ*apiAT2* and Δ*apiAT2* parasites complemented with a constitutively expressed *Tg*ApiAT2 (c*Tg*ApiAT2/Δ*apiAT2*). (C) WT (TATi) and *ΔapiAT3* sub-family mutants. (D) WT (TATi/Tomato) and Δ*apiAT5* sub-family mutants. (E) WT (TATi/Tomato) and Δ*apiAT6* subfamily mutants. (F) Δ*apiAT7* sub-family mutants. Note that the TATi/Tomato strain served as WT strain for the Δ*apiAT2*, Δ*apiAT5*, Δ*apiAT6*, and Δ*apiAT7* sub-family mutants, and the identical image of the TATi/Tomato plaque assay is shown in B, D and E to facilitate interpretation of the data. All images are from the same experiment, and are representative of three independent experiments.

### *Tg*ApiAT5-3 is important for amino acid homeostasis in *T. gondii*

*Tg*ApiAT proteins that are important for parasite growth are likely to have critical roles in nutrient acquisition. In the remainder of this manuscript, we focus on one such protein, *Tg*ApiAT5-3. Our previous study of *Tg*ApiAT1 and *Pb*ApiAT8-1 indicated a key role for these transporters in cationic amino acid uptake (5), and we hypothesized that *Tg*ApiAT5-3 could also function as an amino acid transporter. To investigate this possibility, we incubated WT and Δ*apiAT5-3* parasites in medium containing a mixture of [^13^C]-labelled amino acids for 15 mins. Polar metabolites were extracted from parasite lysates and analyzed by GC-MS. These analyses were used to quantitate the levels of intracellular amino acids and determine the extent of labeling with exogenous amino acids based on [^13^C]-enrichment in each of the two strains. Strikingly, both the abundance and fractional labelling of [^13^C]-L-tyrosine (L-Tyr) was reduced significantly in the Δ*apiAT5-3* strain (Fig 4). Abundance and fractional labeling of a number of other [^13^C]-amino acids were altered significantly in the Δ*apiAT5-3* strain, although none to the same extent as L-Tyr (Fig 4). These data indicate that *Tg*ApiAT5-3 is important for amino acid homeostasis, playing a key role in the uptake of L-Tyr into the parasite.

**Fig 4.**
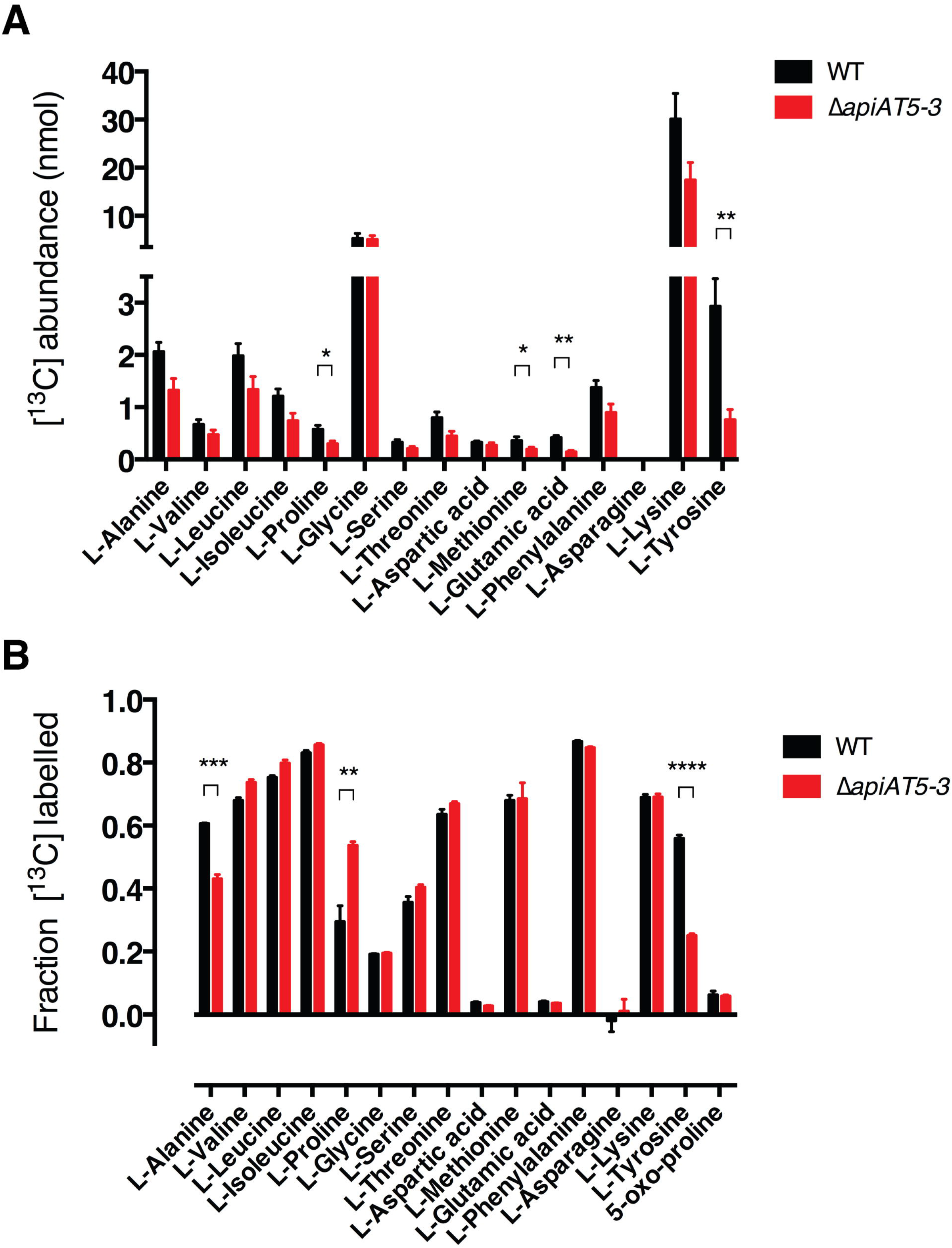
Analysis of [^13^C] amino acid uptake into WT and Δ*apiAT5-3* parasites reveals a role for *Tg*ApiAT5-3 in amino acid homeostasis. (A-B) Extracellular WT or Δ*apiAT5-3* tachyzoites were incubated in medium containing [^13^C]-L-amino acids for 15 min. Polar metabolites were extracted and amino acid abundance (A) and levels of [^13^C]-amino acid enrichment (B) in WT (black) and Δ*apiAT5-3* (red) tachyzoites determined by GC-MS. Only L-amino acids that could be detected in all experiments are shown. The data are averaged from three independent experiments and error bars represent ± s.e.m. (*, P < 0.05; **, P < 0.01; ***, P < 0.001, ****, P < 0.0001; Student’s *t* test. Where significance values are not shown, the differences were not significant; P > 0.05).

### *Tg*ApiAT5-3 is an aromatic and large neutral amino acid uniporter with exchange activity

To characterize the substrate specificity of *Tg*ApiAT5-3 further, we expressed HA-tagged *Tg*ApiAT5-3 in *Xenopus laevis* oocytes, and confirmed its expression and plasma membrane localization by western blotting (Fig S4A). Given the GC-MS data implicating *Tg*ApiAT5-3 in L-Tyr uptake (Fig 4), we hypothesised that *Tg*ApiAT5-3 transports L-Tyr. We compared the uptake of radiolabelled [^14^C]-tyrosine ([^14^C]Tyr) into oocytes expressing *Tg*ApiAT5-3 relative to uninjected oocytes. Under the conditions of the experiment, there was a significant, 7-fold increase in the initial uptake rate of [^14^C]Tyr into oocytes expressing *Tg*ApiAT5-3 compared to uninjected control oocytes (Fig 5A).

**Fig 5.**
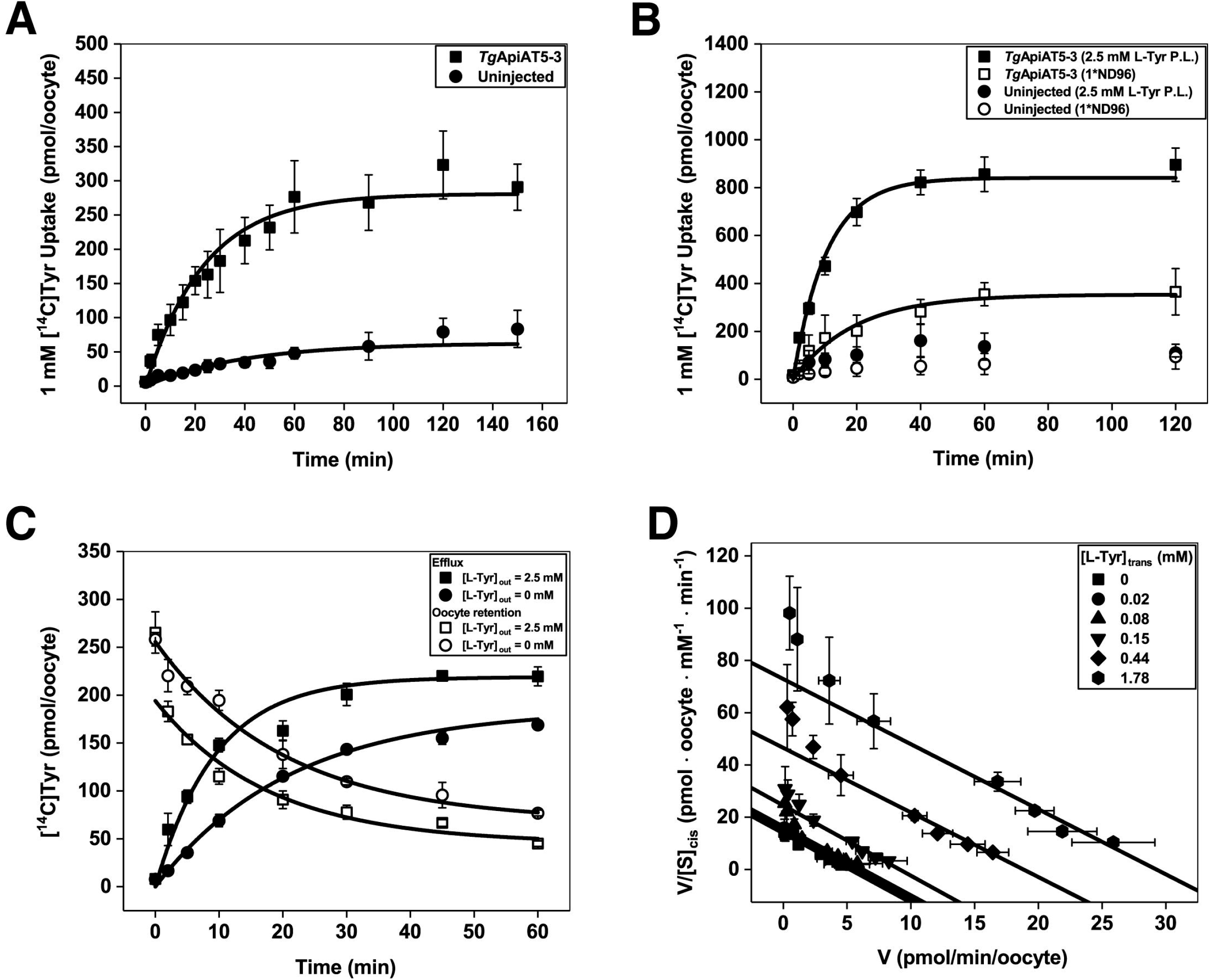
*Tg*ApiAT5-3 is an L-tyrosine transporter that is stimulated by the presence of L-tyrosine on the *trans* side of the membrane. (A) Time course for the uptake of [^14^C]Tyr into *X. laevis* oocytes expressing *Tg*ApiAT5-3 (squares) or into uninjected oocytes (circles). Uptake was measured in the presence of 1 mM L-Tyr. Each data point represents the mean uptake in 10 oocytes from a single experiment ± standard deviation, and the data are representative of 3 independent experiments. A first order rate equation was fitted to each time course (R^2^ = 0.97 for *Tg*ApiAT5-3-expressing oocytes and R^2^ = 0.77 for uninjected controls). Both the rate constant for [^14^C]Tyr uptake and the maximal [^14^C]Tyr uptake measured in *Tg*ApiAT5-3-expressing oocytes were significantly higher than those measured in uninjected oocytes (P < 0.01, Student’s *t* tests). (B) *Tg*ApiAT5-3-expressing oocytes (squares) and uninjected oocytes (circles) were preloaded with L-Tyr by incubation in 2.5 mM L-Tyr (filled symbols) for 32 or 72 hr, respectively, or not preloaded (open symbols). Following the preincubation period, uptake of [^14^C]Tyr was measured in medium containing 1 mM L-Tyr. Data show the mean uptake in 10 oocytes from a single experiment ± standard deviation, and are representative of 3 independent experiments. First order rate equations were fitted to the uptake time courses for the preloaded and non-preloaded *Tg*ApiAT5-3-injected oocytes (R^2^ = 0.98 for preloaded, and R^2^ = 0.95 for non-preloaded oocytes). Both the first order rate constants for [^14^C]Tyr uptake and the maximal [^14^C]Tyr uptake were significantly higher in preloaded compared to nonpreloaded *Tg*ApiAT5-3-expressing oocytes (P < 0.01, Student’s *t* tests). (C) *Tg*ApiAT5-3-expressing oocytes were preloaded by incubation in 1 mM [^14^C]Tyr for 32 hr. Subsequent efflux (filled symbols) and retention (open symbols) of the preloaded labelled substrate was measured over the time-course indicated, in the presence of an extracellular medium containing 2.5 mM L-Tyr (squares) or extracellular medium lacking of L-Tyr (circles). Data show the mean efflux and retention ± standard deviation in 3 replicates (measuring efflux/retention from 5 oocytes each) from a single experiment, and are representative of 3 independent experiments. (D) *Trans*-stimulated initial rate kinetic analysis of L-Tyr transport by *Tg*ApiAT5-3. The rate of L-Tyr uptake was measured at a range of [L-Tyr] concentrations in the external medium (i.e. [L-Tyr]_cis_) in *Tg*ApiAT5-3-expressing oocytes preloaded with 0 mM to 2.5 mM L-Tyr (i.e. [L-Tyr]_trans_). The *Tg*ApiAT5-3-mediated uptake (calculated by subtracting the uptake in uninjected oocytes from the uptake in *Tg*ApiAT5-3-expressing oocytes) at each [L-Tyr]_trans_ condition tested conformed to a Michaelis-Menten kinetic model (R^2^ > 0.90 for all non-linear regressions). The data were fitted to a Scatchard linear regression (0.89 ≤ R^2^ ≤ 0.98 for all linear regressions). Data show the mean uptake rate ± standard deviation in 10 oocytes from a single experiment, and are representative of 2 independent experiments.

While loss of TgApiAT5-3 led to reduced uptake of [^13^C]-tyrosine, levels of ^13^C enrichment in some other amino acids increased (Fig 4). This gave rise to the possibility that *Tg*ApiAT5-3 has exchange activity. To investigate this, we tested whether the uptake of [^14^C]Tyr was stimulated by the presence of amino acids on the *trans* side of the membrane (i.e. *inside* the oocyte). Following preliminary experiments to optimise the preloading of L-Tyr into oocytes (not shown), we measured [^14^C]Tyr uptake over 2 hr at an extracellular concentration of 1 mM L-Tyr in *Tg*ApiAT5-3-injected or uninjected oocytes that had been pre-loaded in medium containing 2.5 mM unlabelled L-Tyr. The initial rate of [^14^C]Tyr uptake in *Tg*ApiAT5-3-expressing oocytes preloaded with L-Tyr was 3-fold higher than in *Tg*ApiAT5-3-expressed oocytes that were not pre-loaded (Fig 5B), indicating that L-Tyr uptake into *Tg*ApiAT5-3-expressing oocytes was stimulated by L-Tyr on the *trans* side of the membrane. We next tested whether L-Tyr efflux was also *trans*-stimulated. We preloaded *Tg*ApiAT5-3-expressing and uninjected oocytes with 1 mM [^14^C]Tyr and measured the efflux and retention of the radiolabel upon the addition of 2.5 mM unlabelled L-Tyr to the external medium. [^14^C]Tyr efflux was increased, and [^14^C]Tyr retention reduced, in *Tg*ApiAT5-3-expressing oocytes exposed to 2.5 mM L-Tyr compared to those in external medium lacking L-Tyr (Fig 5C). Nevertheless, we still observed some [^14^C]Tyr efflux over time in the absence of *trans*-substrate (Fig 5C). We observed no differences in [^14^C]Tyr efflux or retention in control uninjected oocytes upon incubation of oocytes in 2.5 mM L-Tyr compared to incubation in buffer lacking L-Tyr (Fig S4B). These data indicate that the transporter operates more effectively under ‘exchange conditions’ than under conditions in which it is mediating a unidirectional flux.

To examine the exchange activity of *Tg*ApiAT5-3 further, we investigated the kinetic properties of L-Tyr transport in more detail. Steady-state kinetic parameters for exchangers must be conducted at different *trans*- and *cis*-substrate concentrations to determine accurate K_0.5_ values (19). We examined uptake kinetics of [^14^C]Tyr following the preloading of oocytes with different L-Tyr concentrations (note that this, and all subsequent, uptake measurements were conducted over 10 min, representing the approximate initial rate conditions for substrate influx; Fig 5B). Both Michaelis-Menten analysis (not shown) and Scatchard linear regressions (Fig 5D) demonstrated that the apparent K_0.5_ values for L-Tyr uptake into oocytes was unaffected by the cytosolic L-Tyr concentration. The gradients of fitted Scatchard plots are negative reciprocals of the apparent K_0.5_ and, therefore, equivalent slopes reflect similar apparent transport affinities. K_0.5_ values were consistent for both Scatchard and Michaelis-Menten plots derived from the same data, ranging from 0.25 to 0.40 μM (Table 1). As expected for *trans*-stimulated uptake, maximum rate (V_max_) values increased in proportion to the concentration of preloaded L-Tyr (Fig 5D; Table 1).

**Table 1:**
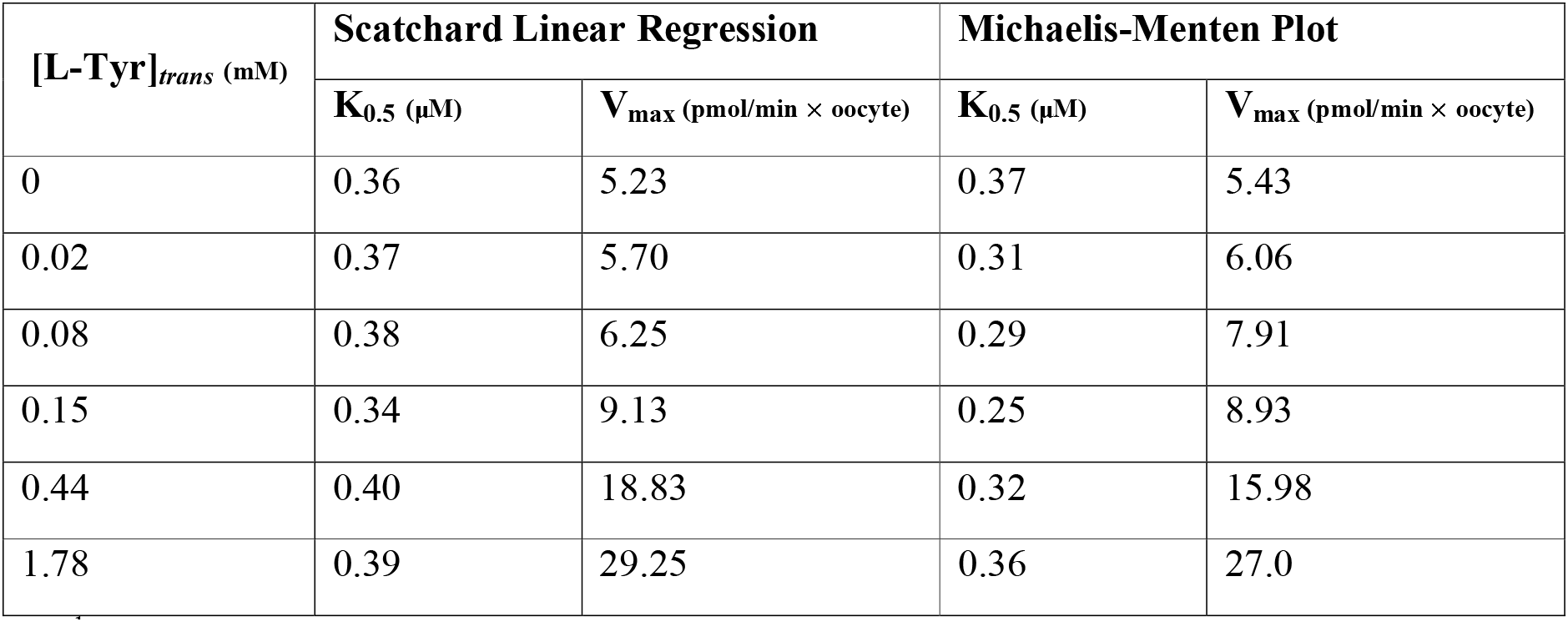
Michaelis-Menten kinetic parameters for the initial rate of L-Tyr uptake by *Tg*ApiAT5-3^1^.

| [L-Tyr]<sub>trans</sub> (mM) | Scatchard Linear Regression |  | Michaelis-Menten Plot |  |
| --- | --- | --- | --- | --- |
|  | K<sub>0.5</sub> (μM) | V<sub>max</sub> (pmol/min × oocyte) | K<sub>0.5</sub> (μM) | V<sub>max</sub> (pmol/min × oocyte) |
| 0 | 0.36 | 5.23 | 0.37 | 5.43 |
| 0.02 | 0.37 | 5.70 | 0.31 | 6.06 |
| 0.08 | 0.38 | 6.25 | 0.29 | 7.91 |
| 0.15 | 0.34 | 9.13 | 0.25 | 8.93 |
| 0.44 | 0.40 | 18.83 | 0.32 | 15.98 |
| 1.78 | 0.39 | 29.25 | 0.36 | 27.0 |

<sup>1</sup> Values are based on the experiment shown in Fig 5D.

^1^Values are based on the experiment shown in Fig 5D.

To assess whether *Tg*ApiAT5-3 activity might function as an ion:amino acid co-transporter, we conducted systematic ion-replacement uptake experiments as described previously (5). We observed no significant change in [^14^C]Tyr uptake under any ion-replacement conditions, suggesting that the transporter does not co-transport any of the ions tested (Fig S4C). To determine whether *Tg*ApiAT5-3 is an electrogenic transporter (as is the case for *Tg*ApiAT1; (5)), we perfused oocytes expressing *Tg*ApiAT5-3 with L-Tyr and measured currents using a two-electrode voltage clamp configuration. We observed no net current movement, including under conditions in which the membrane potential and pH gradient across the membrane were altered (Fig S4D). Together, these data indicate that L-Tyr transport by *Tg*ApiAT5-3 does not co-transport any charged species.

Incubation of WT and Δ*apiAT5-3* parasites in a [^13^C]-amino acid mix resulted in significant increases or decreases in the ^13^C-labelling of several amino acids (Fig 4), suggesting that *Tg*ApiAT5-3 may transport a range of amino acids. To investigate the substrate specificity of *Tg*ApiAT5-3 further, we measured the uptake of 500 μM [^14^C]Tyr in oocytes expressing *Tg*ApiAT5-3 in the presence of equimolar amounts of other unlabelled L-amino acids on the *cis* side of the membrane. Only unlabelled L-tryptophan (L-Trp) resulted in a significant inhibition of L-Tyr uptake under the conditions tested (Fig S5A). To examine the *trans*-stimulated influx specificity of *Tg*ApiAT5-3, we pre-injected mixtures of L-amino acids and other metabolites into oocytes expressing *Tg*ApiAT5-3, then measured uptake of [^14^C]Tyr. *Tg*ApiAT5-3-mediated [^14^C]Tyr uptake was *trans*-stimulated by pre-injected mixtures of L-amino acids, but not amino acid derivatives, D-amino acids, nucleotides, nitrogenous bases, or sugars (Fig S5B; Table S2). To identify which amino acids might be responsible for this *trans*-stimulation, we systematically pre-injected oocytes expressing *Tg*ApiAT5-3 with 5 mM of each of the 20 proteinogenic amino acids in the mixture (with the exception of L-Tyr which was preloaded to equilibrium), before measuring *trans*-stimulated uptake of [^14^C]Tyr. We found that aromatic and large neutral amino acids, but not smaller or charged amino acids, *trans*-stimulated [^14^C]Tyr uptake (Fig 6A).

**Fig 6.**
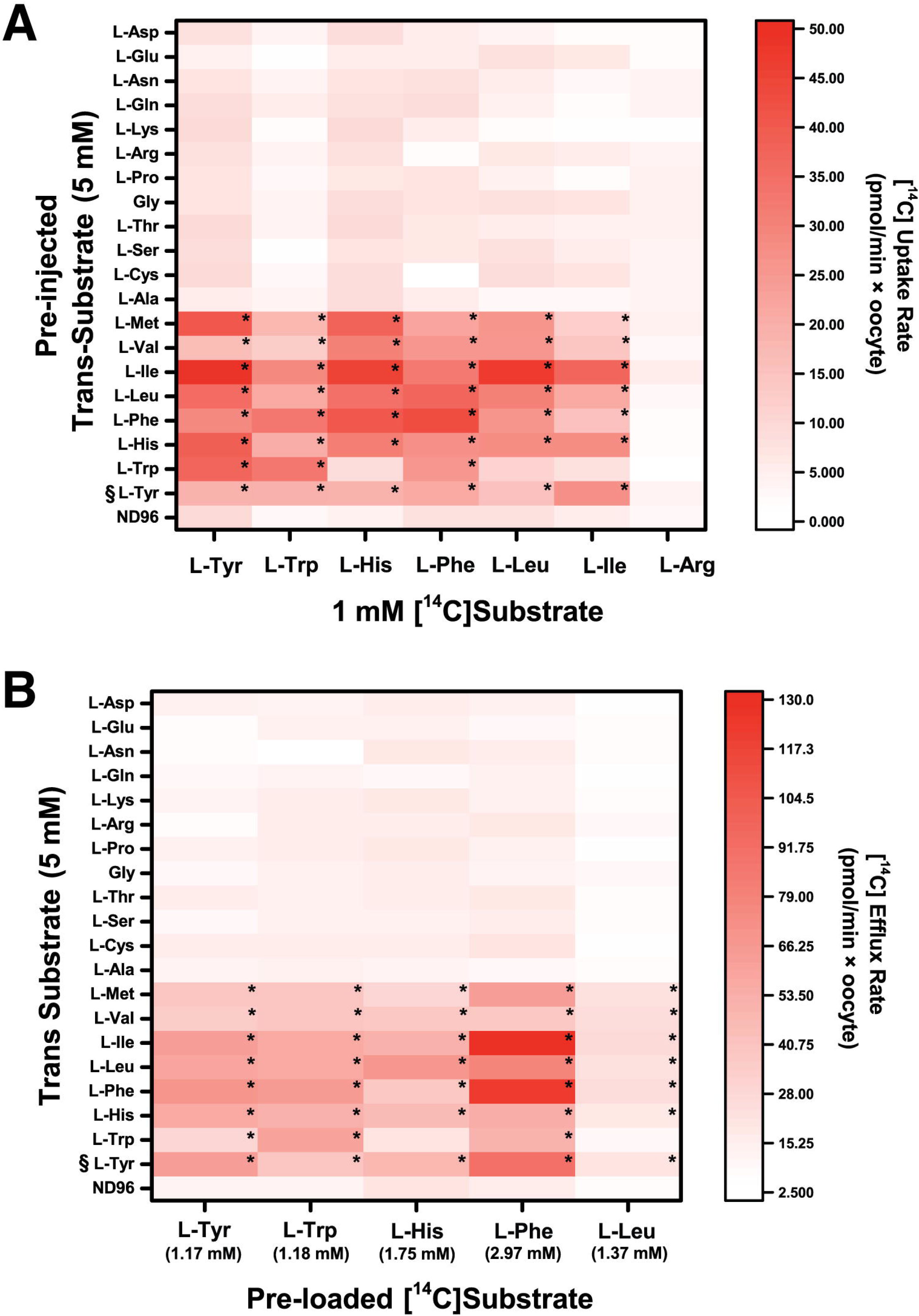
*Tg*ApiAT5-3 is an exchanger for aromatic and large neutral amino acids. (A) *Tg*ApiAT5-3-expressing oocytes were pre-injected with a range of L-amino acids at a calculated oocyte cytosolic concentration of 5 mM, with the exception of L-Tyr (§) which was preloaded via incubation in 2.5 mM L-Tyr for 32 hr, or were not pre-injected (ND96). Subsequent uptake of [^14^C]-labelled amino acids was measured over 10 minutes and normalised to uptake per minute. Each box in the heat map shows the mean uptake in 10 oocytes from a single experiment, representative of 3 independent experiments. The statistical analyses compare pre-injected/pre-loaded oocytes to ND96 controls for each substrate tested (*, P < 0.05, one-way ANOVA, Dunnet’s post-hoc test. Where significance values are not shown, the differences are not significant, P > 0.05). (B) *Tg*ApiAT5-3-expressing oocytes were preloaded with a range of [^14^C]-labelled amino acids (calculated final concentrations shown beneath each substrate), and efflux of these substrates was measured over 5 min in the absence of external amino acids (ND96) or in the presence of 5 mM external amino acids (with the exception of L-Tyr (§), which was present at a concentration of 2.5 mM), and normalised to efflux per minute. Each box in the heat map shows the mean efflux from 3 replicates (each comprised of 5 oocytes) from a single experiment, representative of 3 independent experiments. Statistical analyses compare trans substrates to ND96 controls for each efflux substrate tested (*, P < 0.05, one-way ANOVA, Dunnet’s post-hoc test. Where significance values are not shown, the differences are not significant, P > 0.05).

We tested the uptake of a range of [^14^C]-labelled aromatic and large neutral amino acids, including L-Trp, L-histidine (L-His), L-phenylalanine (L-Phe), L-leucine (L-Leu) and L-isoleucine (L-Ile) into oocytes expressing *Tg*ApiAT5-3. In the absence of a trans substrate, the rate of uptake of aromatic and large neutral amino acids were significantly increased compared to uninjected oocytes (1.7-fold increase for L-Trp, 3.1-fold increase for L-His, 5.5-fold increase for L-Phe, 2.2-fold increase for L-Leu, and 2.4-fold increase for L-Ile; Fig S5C). The uptake of the cationic amino acid L-Arg did not differ between *Tg*ApiAT5-3 injected and uninjected oocytes (Fig S5C). As was seen for L-Tyr, uptake of the tested aromatic and large neutral amino acids was *trans*-stimulated by aromatic and large neutral amino acids, and not by smaller or charged L-amino acids (Fig 6A). The uptake of L-Arg was not *trans*-stimulated by any of the amino acids tested (Fig 6A). We next measured efflux of preloaded [^14^C]-labelled L-Tyr, L-Trp, L-His, L-Phe and L-Leu in the presence of all 20 proteinogenic amino acids in the extracellular medium. We observed the same substrate specificity, with aromatic and large neutral L-amino acids stimulating the efflux of the [^14^C]-labelled substrates tested (Fig 6B).

From the experiments conducted in this section, we conclude that *Tg*ApiAT5-3 can mediate the transport of aromatic and large neutral amino acids. The transporter can function as a uniporter, but has a strong propensity for exchange, implying a role for *Tg*ApiAT5-3 in the homeostasis of a range of aromatic and large neutral amino acids.

### *Tg*ApiAT5-3 is important for tyrosine uptake into parasites

The [^13^C] amino acid uptake data and oocyte experiments indicate a role for *Tg*ApiAT5-3 in L-Tyr uptake into the parasite (Fig 4; Fig 5). To test the importance of *Tg*ApiAT5-3 for L-Tyr uptake in *T. gondii*, we measured the kinetics of [^14^C]Tyr uptake in WT and Δ*apiAT5-3* parasites. The initial rate of [^14^C]Tyr uptake in Δ*apiAT5-3* parasites was decreased by 8.5-fold compared to that in WT parasites (Fig 7A; Fig S6A). Both [^14^C]Tyr uptake and parasite growth were increased upon complementation of the Δ*apiAT5-3* mutant with a constitutively expressed copy of *Tg*ApiAT5-3 (c*Tg*ApiAT5-3/Δ*apiAT5-3*; Fig 7A; Fig S7). However, uptake was not restored to WT levels, perhaps as a result of different levels of expression of *Tg*ApiAT5-3 in the WT and complemented strains.

**Fig 7.**
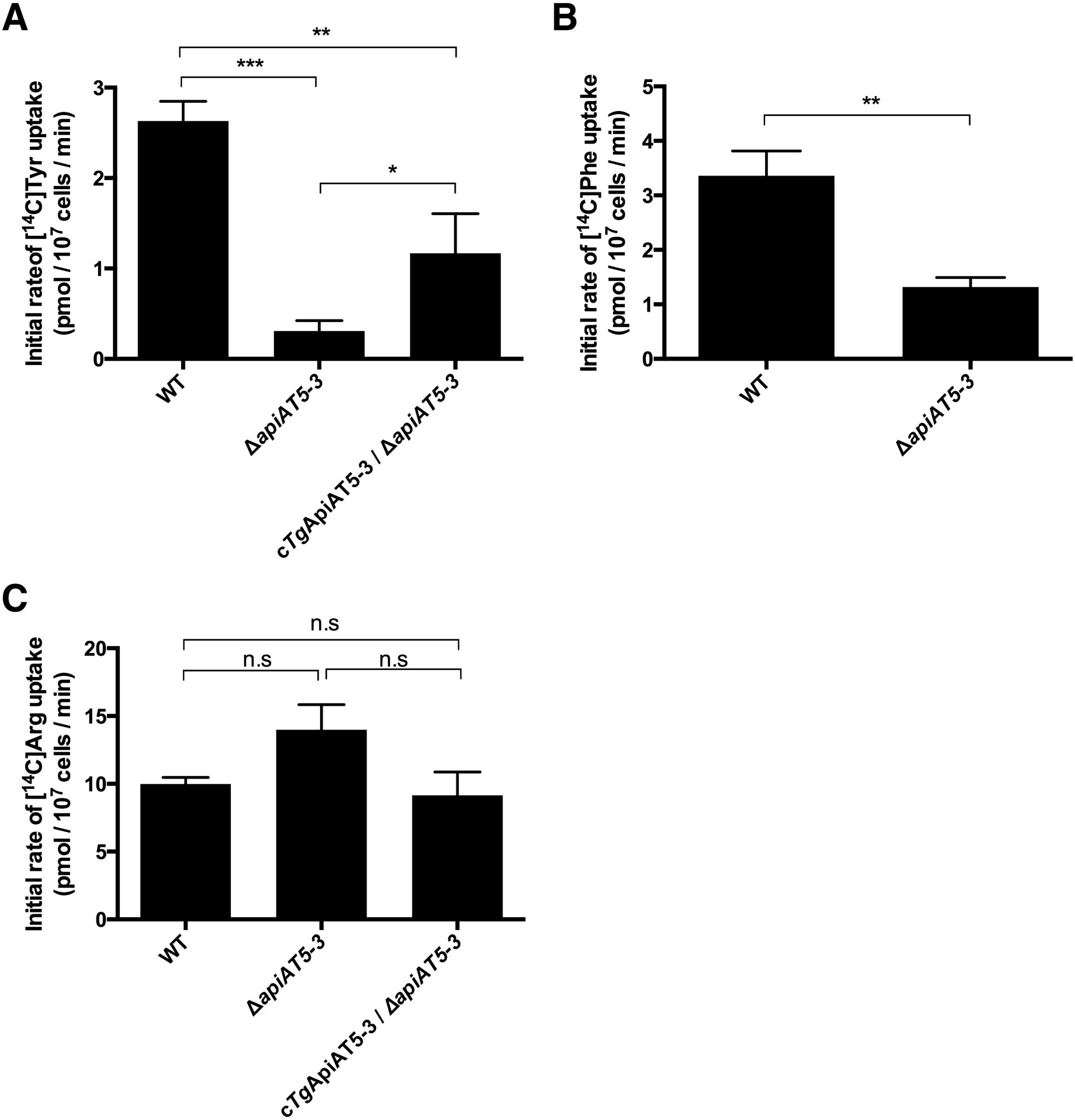
*Tg*ApiAT5-3 mediates the uptake of L-tyrosine and L-phenylalanine into *T. gondii*. Initial rate of uptake of (A) [^14^C]Tyr, (B) [^14^C]Phe, and (C) [^14^C]Arg, in WT, Δ*apiAT5-3*, and (for A and C) c*Tg*ApiAT5-3/Δ*apiAT5-3* strain parasites. Uptake was measured in PBS-glucose containing either 60 μM unlabelled L-Tyr and 0.1 μCi/mL [^14^C]Tyr (A), 15 μM unlabelled L-Phe and 0.1 μCi/mL [^14^C]Phe (B), or 100 μM unlabelled L-Arg and 0.1 μCi/mL [^14^C]Arg (C). The initial rates of transport for each substrate were computed from the initial slopes of the fitted single-order exponential curves (Fig S6), and represent the mean ± SEM from three independent experiments (* *P*<0,05; ** *P*<0.01; *** *P*<0.001; n.s. = not significant; Student’s *t* test).

Our previous data indicated that *Tg*ApiAT5-3 can transport other aromatic amino acids (Fig 6, Fig S5A). We measured uptake of [^14^C]Phe in WT and Δ*apiAT5-3* parasites. We found a significant, 2.5-fold decrease in the initial rate of [^14^C]Phe uptake in parasites lacking *Tg*ApiAT5-3 (Fig. 7B; Fig S6B). We were unable to detect robust levels of uptake of [^14^C]Trp in either WT or Δ*apiAT5-3* parasites, precluding analysis of the role of *Tg*ApiAT5-3 in uptake of this amino acid into parasites. As a control, we measured uptake of [^14^C]Arg, which is not transported by *Tg*ApiAT5-3 (Fig 6; Fig S5A), in WT, Δ*apiAT5-3* and c*Tg*ApiAT5-3/Δ*apiAAT5-3* strain parasites. We found that the rate of [^14^C]Arg uptake did not differ significantly between these parasite lines (Fig 7C; Fig S6C), indicating that the defect we observe in the uptake of aromatic amino acids is specific, and does not represent a general defect in amino acid uptake.

### Growth of parasites lacking *Tg*ApiAT5-3 is modulated by the concentrations of aromatic amino acids in the growth medium

We next investigated the dependence of the growth of Δ*apiAT5-3* parasites on the concentration of L-Tyr in the culture medium. WT and Δ*apiAT5-3* parasites were grown in DMEM containing 0 – 2.5 mM L-Tyr. Growth of WT parasites in the absence of L-Tyr was severely impaired (Fig S8A), consistent with a previous study that indicated *T. gondii* parasites are auxotrophic for this amino acid (20). WT parasites grew normally in [L-Tyr] as low as 10 μM (Fig 8A). By comparison, growth of Δ*apiAT5-3* parasites was negligible at [L-Tyr] of 156 μM and below, and severely impaired at concentrations below 1 mM (Fig 8A). We also measured growth of WT, Δ*apiAT5-3* and c*TA*piAT5-3/Δ*apiAT5-3* parasites grown in DMEM vs DMEM containing 2.5 mM L-Tyr, by plaque assay. Consistent with the results of the fluorescence growth assays, plaque assays revealed that impaired growth of Δ*apiAT5-3* parasites was restored by growth in 2.5 mM L-Tyr (Fig S7).

**Fig 8.**
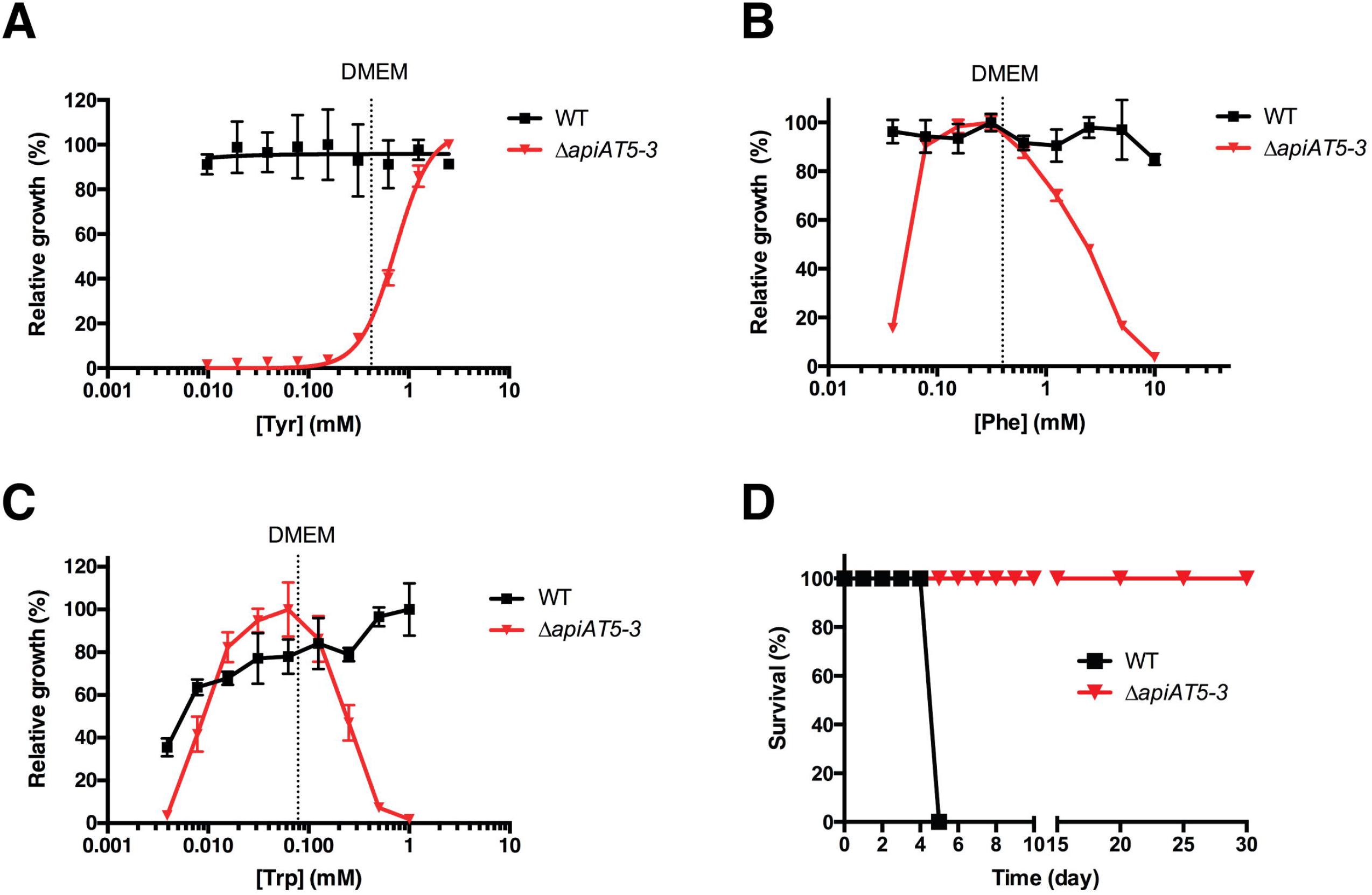
*in vitro* growth of parasites lacking *Tg*ApiAT5-3 is modulated by the concentration of aromatic amino acids in the growth medium, and *Tg*ApiAT5-3 is important for parasite virulence. (A-C) Fluorescence growth assay for WT (black) and Δ*apiAT5-3* (red) parasites cultured for 5 days in DMEM containing a range of L-Tyr (A), L-Phe (B), or L-Trp (C) concentrations. Growth is expressed as a percentage of maximum growth measured on day 5 for each parasite strain. For Δ*apiAT5-3* in (A), a sigmoidal curve has been fitted to the data. All data shown are averaged from three technical replicates (mean ± standard deviation), and are representative of those obtained in three independent experiments. The concentration of the respective L-amino acids in standard DMEM are depicted by a dashed line. (D) Five Balb/c mice were infected intraperitoneally with 1,000 RHΔ*hxgprt*/Tomato (black) or Δ*apiAT5-3* in RHΔ*hxgprt*/Γomato (red) strain parasites and monitored for symptoms of toxoplasmosis.

These data indicate that uptake of L-Tyr via *Tg*ApiAT5-3 is important for the growth of *T. gondii* parasites in standard *in vitro* conditions. They also point to the existence of an alternative L-Tyr uptake pathway that can mediate sufficient L-Tyr uptake for parasite growth when parasites are cultured in medium containing ≥ 1 mM L-Tyr.

To assess the physiological significance of *Tg*ApiAT5-3-mediated L-Phe uptake for parasite growth, WT and Δ*apiAT5-3* were cultivated in medium containing 0 to 10 mM L-Phe (at a constant [L-Tyr] of 423 μM, the concentration of this amino acid in DMEM). WT parasites grew minimally in the absence of L-Phe (Fig S8B), indicating that *T. gondii* is auxotrophic for this amino acid, but grew normally at [L-Phe] of 39 μM and above (Fig 8B). By contrast, growth of Δ*apiAT5-3*parasites was impaired at 39 μM [L-Phe] (Fig 8B). Δ*apiAT5-3*parasites grew optimally at [L-Phe] between 78 μM and 625 μM, but parasite growth decreased at [L-Phe] of 1.25 mM and above (Fig 8B).

We next measured the growth of WT and Δ*apiAT5-3* parasites in medium containing 0 – 1 mM L-Trp (and a constant [L-Tyr] of 423 μM). WT parasites grew minimally in the absence of L-Trp (Fig S8C), consistent with a previous study that indicated *T. gondii* is auxotrophic for this amino acid (21). WT parasites grew optimally at 1 mM [L-Trp], and exhibited decreased growth with decreasing [L-Trp] (Fig 8C). By contrast, Δ*apiAT5-3* parasites grew optimally at 16 – 125 μM L-Trp (Fig 8C). Growth of Δ*apiAT5-3* parasites was negligible at 4 μM L-Trp (a concentration at which growth of WT parasites is only moderately impaired), and also decreased at [L-Trp] of 250 μM and above (Fig 8C), mirroring the effects observed with growth of the mutant at a range of [L-Phe].

Together, these data reveal that *Tg*ApiAT5-3 is required for parasite growth at low exogenous concentrations of L-Phe and L-Trp, but not at intermediate concentrations. This points to the existence of other L-Phe and L-Trp uptake pathways in the parasite. Notably, *Tg*ApiAT5-3 is required for parasite growth at high exogenous L-Phe and L-Trp concentrations. This observation suggests that high concentrations of L-Phe or L-Trp may competitively inhibit uptake of L-Tyr via the alternative, non-*Tg*ApiAT5-3 L-Tyr uptake pathway (considered further in the Discussion).

### *Tg*ApiAT5-3 is important for parasite virulence

Standard media formulations used to culture parasite *in vitro* do not necessarily reflect the amino acid concentrations that parasites encounter *in vivo*. Since the importance of *Tg*ApiAT5-3 for parasite growth *in vitro* is dependent on the concentrations of aromatic amino acids in the growth medium (Fig 8A-C), we investigated the importance of *Tg*ApiAT5-3 for parasite virulence in a mouse infection model. In preliminary experiments, we found that the parental TATi/Tomato WT parasite strain was avirulent in mice, precluding an analysis of whether Δ*apiAT5-3* parasites are virulent. We therefore remade the Δ*apiAT5-3* mutant in virulent RHΔ*hxgprt*/Tomato strain parasites (22). Δ*apiAT5-3*/RHΔ*hxgprt*/Tomato parasites were defective in uptake of [^14^C]Tyr (Fig S9A), and were dependent on high levels of exogenous L-Tyr for growth (Fig S9B). Notably, when we compared the growth of Δ*apiAT5-3*/RHΔ*hxgprt*/Tomato parasites to parental RHΔ*hxgprt*/Tomato parasites by plaque assay we found that, although growth in 2.5 mM Tyr partially restored growth of Δ*apiAT5-3*/RHΔ*hxgprt*/Tomato strain parasites, this strain still grew slower than the parental parasites (Fig S9C). This is in contrast to the equivalent experiment with Δ*apiAT5-3*/TATi/Tomato parasites (Fig S7), and suggests some inter-strain differences in the importance of *Tg*ApiAT5-3 for *in vitro* growth, or in the availability of L-Tyr in the intracellular niche created by different strains.

We infected BALB/c mice intraperitoneally with 10^3^ WT (RHΔ*hxgprt*/Tomato) or Δ*apiAT5-3*/RHΔ*hxgprt*/Tomato parasites and monitored disease progression. Mice infected with WT parasites exhibited symptoms of toxoplasmosis and were euthanized 6 days post-infection (Fig 8D). In contrast, mice infected with Δ*apiAT5-3* parasites exhibited no symptoms of toxoplasmosis across the entire 62 days of the experiment (Fig 8D), indicating that *Tg*ApiAT5-3 is essential for parasite virulence.

## Discussion

The evolution of apicomplexan ancestors from a free-living to an obligate parasitic life-style was associated with the loss of numerous biosynthetic pathways, including those for amino acid synthesis (23, 24). In particular, *T. gondii* is auxotrophic for many amino acids, including the aromatic amino acids L-Tyr, L-Phe and L-Trp ((5, 20, 21, 25–27); this study). Parasites must scavenge essential amino acids from their environment, although, in the case of apicomplexan parasites, how they do so is poorly understood. Here, we characterise a family of proteins predicted to function as transporters in apicomplexans. Using a combination of *in vitro, in vivo*, and heterologous expression studies, we provide evidence that these transporters localize to the parasite plasma membrane, and show that *Tg*ApiAT5-3 functions as an aromatic amino acid transporter in *T. gondii*. We have previously demonstrated that other members of this family, which we now call *Tg*ApiAT1 (previously *Tg*NPT1) and *Pb*ApiAT8-1 (previously *Pb*NPT1), transport cationic amino acids (5). Our phylogenetic analyses indicate that ApiAT proteins are found throughout the apicomplexan phylum. To reflect these functions and phylogenetic affinities, we propose to rename this protein family the *Api*complexan *A*mino acid *T*ransporters (ApiATs).

The ApiAT protein family is found in all apicomplexan species that we analyzed, as well as in chromerids, which are free-living relatives of apicomplexans. Chromerids are prototrophic for amino acids, although growth of these algae can be enhanced by the addition of glutamate and glycine to the growth medium, suggesting they can also acquire exogenous amino acids (28). Understanding the localization and function of chromerid ApiATs may provide valuable clues to the evolution of this transporter family in apicomplexans. The similarity of ApiATs to mammalian LAT3/4-type amino acid transporters suggests that the ancestral function of ApiATs was amino acid transport. The ApiATs appear to have undergone expansion in various apicomplexan lineages, including *Plasmodium* spp, *T. gondii* and piroplasms such as *Babesia* spp and *Theileria* spp, whereas only a single representative is present in *Cryptosporidium* spp, an early-diverging lineage of apicomplexans (29). These observations are consistent with ancestral apicomplexans containing a single ApiAT protein that diversified in various lineages of the phylum to encompass new and/or more selective amino acid substrate selectivities.

A similar expansion has been observed in the amino acid/auxin permease (AAAP) family of trypanosomatid parasites, in which fourteen AAAP genes arose from a single AAAP gene locus through a series of gene duplication events in ancestral trypanosomatids (30). AAAP expansion is likely to reflect an early parasitic innovation that contributed to establishing parasite dependency on the host organism, and thereby facilitating the evolution of parasitism in trypanosomatids (23, 30). By contrast, much of the expansion in the ApiAT family appears to have occurred subsequent to the diversification of the major lineages in the phylum. Of the nine ApiAT subfamilies that we define, only the ApiAT2 subfamily is broadly distributed amongst the major apicomplexan lineages (Fig 1), suggesting its presence before these lineages diverged. Several subfamilies have undergone expansion within lineages. For example, the ApiAT3, 5, 6 and 7 subfamilies contain multiple members within coccidians (*T. gondii, N. caninum* and *E. tenella*), while piroplasms contain multiple ApiAT2 subfamily proteins (Fig 1).

Much of the expansion of ApiAT proteins, then, appears to have occurred subsequent to the evolution of parasitism in this phylum. An intriguing possibility is that expansion within ApiAT subfamilies is linked to expansion of these parasites into different hosts, and cell types within those hosts. Across their life cycles, apicomplexans such as *T. gondi, Plasmodium* spp and piroplasms must infect different hosts and/or different cell types with those hosts, which may necessitate amino acid transporters with different substrate affinities and specificities Our data indicate that ten of the sixteen ApiAT family proteins in *T. gondii* are expressed in the tachyzoite stage of the life cycle (Fig 2; (5)). Of the ApiAT5 subfamily, we could only detect expression of *Tg*ApiAT5-3 in tachyzoites (Fig 2), and only *Tg*ApiAT5-3 is important for growth of the tachyzoite stage (Fig 3). This raises the possibility that other *Tg*ApiAT5 subfamily proteins are expressed, and function, at other stages of the life cycle. Interestingly, proteomic studies identified *Tg*ApiAT5-5 in the oocyst proteome (www.toxodb.org), and it could be that this transporter has particular importance at this stage of the parasite life cycle.

Of the fifteen ApiAT family proteins that we were able to disrupt genetically, only the *Tg*ApiAT1, *Tg*ApiAT2 and *Tg*ApiAT5-3 mutants exhibited defects in tachyzoite growth (Fig 3). This corresponds to results from a recent genome-wide CRISPR-based screen, in which these three *Tg*ApiAT family proteins all had low ‘phenotype scores’ (scores between −3.91 and −4.73), an indicator of a gene’s importance for *in vitro* growth of tachyzoites ((31); scores below −1.8 are considered to be indicative of a gene being ‘important’ for parasite growth). The remaining *Tg*ApiAT family proteins had phenotype scores > −0.93 (31), consistent with the results of our targeted knockout approach which indicated that these proteins are not important for parasite growth *in vitro* (Fig 3). We were unable to disrupt the reading frame of *Tg*ApiAT6-1, despite multiple attempts using a guide RNA that targets the *Tg*ApiAT6-1 locus (Table S1). *Tg*ApiAT6-1 has a phenotype score of −5.4 (31). It is likely, then, that our inability to generate a *Tg*ApiAT6-1 knockout is because it is essential for parasite growth.

Our studies of *Tg*ApiAT5-3 demonstrate that this protein is important for parasite growth. We demonstrated that *Tg*ApiAT5-3 is a high affinity L-Tyr uniporter (Fig 5; K_0.5_ ∼0.3 μM, Table 1) and that loss of *Tg*ApiAT5-3 leads to defects in L-Tyr uptake into parasites (Fig 4, Fig 7A). Furthermore, parasites lacking *Tg*ApiAT5-3 were avirulent in mice (Fig 8D), and could only grow at extracellular L-Tyr concentrations above ∼1 mM (Fig 8A), well above the plasma concentration of L-Tyr in mammals (estimated to be 55-90 μM in human plasma and 50-70 μM in mouse plasma; (32, 33)). Together, our data point to an essential role for *Tg*ApiAT5-3 in scavenging L-Tyr from the host.

Our oocyte studies indicated that *Tg*ApiAT5-3 can function as an exchanger, with the rate of uptake of L-Tyr and other aromatic and large neutral amino acids enhanced when equivalent amino acids were present on the *trans* side of the membrane (Fig 6). Mammalian amino acid transporters function either as ‘loaders’ (i.e. uniporters that facilitate the uptake of amino acids into cells) or ‘harmonizers’ (i.e. exchangers that are essential for the maintenance of homeostatic amino acid concentrations) (34). By contrast, our data indicate that *Tg*ApiAT5-3 performs an unusual dual function in facilitating both the net uptake of L-Tyr into the parasite, and maintaining intracellular pools of aromatic and large neutral amino acids through exchange. Maintaining amino acid homeostasis is critical for facilitating cellular metabolism and growth, and it is likely that *Tg*ApiAT5-3 has a critical role in balancing the intracellular concentrations of aromatic and large neutral amino acids in the parasite.

*X. laevis* expression studies revealed that *Tg*ApiAT5-3 transports L-Phe and L-Trp, as well as large neutral amino acids such as L-Leu (Fig 6). This raises the possibility that *Tg*ApiAT5-3 also functions in the net uptake of these amino acids in the parasite. We saw no differences in the fractional labelling of [^13^C]-labelled L-Leu or L-Ile in Δ*apiAT5-3* parasites compared to WT parasites (Fig 4), implying that the uptake of these branched-chain amino acids is facilitated by other transporters in the parasite. Similarly, we observed no defect in the fractional labelling of [^13^C]-labelled L-Phe in parasites lacking *Tg*ApiAT5-3 (Fig 4), although we did observe a defect in the uptake of [^14^C]Phe in the mutant strain (Fig 7C). A possible explanation for this discrepancy lies in the different uptake conditions for these experiments. The [^13^C]-labelled amino acid uptake experiments were performed in medium containing a complex mix of amino acids, whereas [^14^C]Phe uptake was performed in medium containing L-Phe as the sole amino acid. Amino acids such as L-Tyr in the amino acid mix are likely to compete with L-Phe for uptake by *Tg*ApiAT5-3 in WT parasites. Notably, uptake of [^14^C]Tyr into oocytes expressing *Tg*ApiAT5-3 was not impaired by the addition of equimolar amounts of unlabelled L-Phe (Fig S5A), indicating that *Tg*ApiAT5-3 has a greater affinity for L-Tyr than L-Phe. If a transporter’s affinity for L-Tyr is much greater than that for L-Phe, uptake of the latter will be minimal in conditions where the amino acids are present at similar concentrations (as we observed in the oocyte experiments, and as appears to be the case in mammalian cells (32)). This is consistent with the hypothesis that *Tg*ApiAT5-3 plays little role in L-Phe uptake in the parasite. Instead, L-Phe is likely to be taken up via alternative transport pathways (Fig 9).

**Fig 9.**
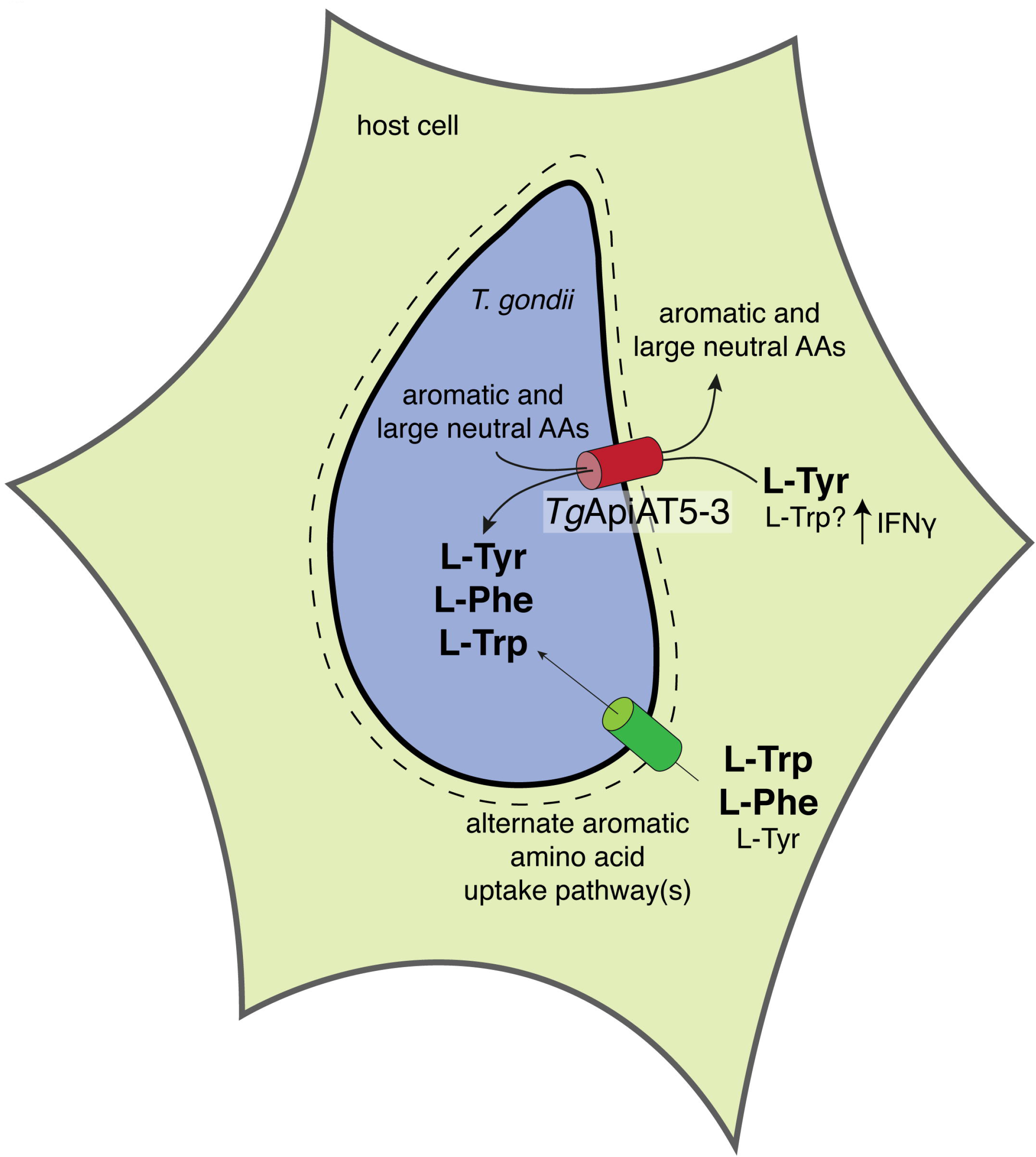
Model for the uptake of aromatic amino acids in *T. gondii*. Depiction of a *T. gondii* parasite (blue) inside a host cell. Aromatic amino acids, including L-Tyr, L-Phe and L-Trp, and large neutral amino acids are thought to be translocated across the parasitophorous vacuole membrane surrounding the parasite (dashed line) through non-selective channels (57). *Tg*ApiAT5-3 (red cylinder) functions as the major L-Tyr uptake pathway in *T. gondii*, and may also have a role in L-Trp uptake following the IFNγ-mediated depletion of this amino acid in the serum. Additionally, *Tg*ApiAT5-3 functions as an exchanger, exporting aromatic and large neutral amino acids from the parasite, and thereby contributing to the homeostasis of these amino acids. The uptake of L-Phe and L-Trp is primarily mediated by alternate, and as yet undefined, uptake pathways (green cylinder). These alternate pathways can mediate sufficient L-Tyr uptake for parasite growth in the absence of *Tg*ApiAT5-3 at high L-Tyr concentrations (when L-Phe and L-Trp concentrations are not correspondingly high).

The importance of *Tg*ApiAT5-3 for L-Trp uptake in *T. gondii* is less clear. We were unable to detect the uptake of either [^13^C]Trp or [^14^C]Trp in parasites under different experimental conditions. In oocyte experiments, L-Trp can effectively out-compete L-Tyr for uptake via *Tg*ApiAT5-3 when present in equimolar amounts (Fig S5A), suggesting that *Tg*ApiAT5-3 has similar affinities for L-Trp and L-Tyr. We observed a defect in growth of Δ*apiAT5-3* parasites at low concentrations of L-Trp in the growth medium (below ∼16 μM; Fig 8C), suggesting that *Tg*ApiAT5-3 may have a role in the uptake of L-Trp at low concentrations. The estimated plasma concentration of L-Trp in humans and mice is ∼60 μM (33), a concentration at which Δ*apiAT5-3* parasites grow optimally *in vitro* (Fig 8C). However, *T. gondii* infection leads to an interferon γ (IFNγ) response in the host organism, which activates the L-Trp-degrading enzyme indoleamine 2,3-dioxygenase, leading to lowered serum levels of L-Trp and decreased parasite growth (21, 35). *Tg*ApiAT5-3 may, therefore, become important for L-Trp uptake following the IFNγ response upon parasite infection *in vivo* (Fig 9). Examining the importance of *Tg*ApiAT5-3 for L-Trp uptake upon IFNγ stimulation will be of particular interest for understanding the interplay between transporter function and the host response to parasite infection.

Δ*apiAT5-3* parasites are capable of growth when the concentration of L-Tyr in the growth medium is ≥1 mM (Fig 8A). This indicates the existence of an alternative L-Tyr uptake pathway that takes up sufficient L-Tyr to enable parasite growth when L-Tyr levels are high (Fig 9). By contrast, Δ*apiAT5-3* parasites grew normally at intermediate concentrations of L-Phe (78 – 625 μM) and L-Trp (31 – 250 μM), but exhibited a dramatic decrease in growth at higher concentrations of both these amino acids (Fig 8B-C). These observations are consistent with the hypothesis that the alternative L-Tyr uptake pathway(s) also functions in L-Phe and L-Trp uptake (Fig 9). At high concentrations of L-Phe and L-Trp, uptake of L-Tyr by this alternative pathway(s) is inhibited by competition with the other aromatic amino acids, preventing parasite growth in the absence of *Tg*ApiAT5-3. This mirrors similar observations in our previous study on the alternative L-Arg uptake pathway in the parasite, in which we showed that high levels of other cationic amino acids inhibit parasite growth in the absence of the selective L-Arg transporter (5).

We propose a model whereby the uptake of L-Tyr into *T. gondii* parasites is mediated primarily by *Tg*ApiAT5-3 (Fig 9). The uptake of L-Phe and L-Trp is mediated primarily by one or more alternative aromatic amino acid transporters. These transporters can also transport L-Tyr, and compensate for loss of *Tg*ApiAT5-3 when L-Tyr levels are high and corresponding levels of L-Phe and L-Trp are lower (Fig 9). *Tg*ApiAT5-3 has an additional role in facilitating homeostasis of these aromatic amino acids and large neutral amino acids (Fig 9).

This model is based on our observations of tachyzoite stage parasites. A recent study examined *T. gondii* aromatic amino acid hydroxylase enzymes, which can interconvert L-Phe and L-Tyr (36). Knockout of these enzymes revealed that they are not required for tachyzoite growth, but are particularly important for producing oocysts following the sexual stages of the parasite life cycle that occur in the felid hosts (36). Given the differences in aromatic amino acid metabolism between tachyzoite and oocyst stages of the life cycle, it is likely that the nature and requirements for aromatic amino acid transporters differs across the life cycle of the parasite. Future studies that investigate the expression and importance of *Tg*ApiAT5-3 and other aromatic amino acid transporters (perhaps other members of the *Tg*ApiAT5 family) across the entire life cycle will be of particular interest.

In this manuscript, we describe an apicomplexan-specific family of plasma membrane transporters that appear to be primarily involved in amino acid uptake. Our findings highlight the evolutionary novelties that must arise to enable parasites to scavenge essential nutrients from their hosts, and also highlight the importance of amino acid scavenging for the growth and virulence of the disease-causing tachyzoite stage *T. gondii*. Future studies that examine the role of other members of this transporter family across the entire life cycle of the parasite will facilitate a better understanding of how these parasites acquire amino acids from their hosts.

## Materials and Methods

### Phylogenetic analysis of the ApiAT protein family

Reciprocal protein BLAST searches in www.eupathdb.org were used to identify orthologues of the five previously identified *Plasmodium falciparum* ApiAT genes in the genomes of the *apicompelxans Plasmodium berghei, Toxoplasma gondii, Cryptosporidium parvum, Eimeria tenella, Neospora caninum, Babesia bovis* and *Theileria annulata*, and the chromerids *Chromera velia* and *Vitrella brassicaformis*. Gene IDs are listed in Table S3.

The sequences of the 67 identified ApiAT proteins were aligned using ClustalX 2.1. The multiple sequence alignment was edited in Jalview (www.jalview.org) to remove poorly aligned blocks. After sequence editing, 452 residues were left for subsequent phylogenetic analysis using PHYLIP v3.69 (evolution.genetics.washington.edu/phylip/getme.html) as described previously (37). Briefly, a consensus maximum likelihood tree and bootstrap values were generated by running the alignment file through the ‘seqboot’ program, which was used to generate 1000 pseudosamples of the alignment. Next, multiple phylogenetic trees were generated from the pseudosamples using the ‘proml’ tree algorithm, using a randomised order of entry and three jumbles. Finally, the multiple phylogenetic trees were converted to a consensus tree with bootstrap values using the program ‘consense’. Trees were viewed using the program FigTree (http://tree.bio.ed.ac.uk/software/figtree/) and annotated using Inkscape (https://inkscape.org/en). For visual representation of the alignment, shading for sequence identity was carried out using the TexShade package for LaTex, using similarity mode with the ‘\fingerprint’ command (https://ctan.org/pkg/texshade?lang=en).

### Parasite culture

Parasites were maintained in human foreskin fibroblasts (HFFs; a kind gift from Holger Schülter, Peter MacCallum Cancer Centre) cultured at 37°C in a humidified 5 % CO_2_ incubator. Unless otherwise noted, parasites were cultured in Dulbecco’s modified Eagle’s medium (DMEM) supplemented with 1 % (v/v) fetal calf serum and antibiotics. ‘Homemade’ media were generated as described previously (5), with amino acids at the concentrations found in DMEM, or modified as specified in the text. Δ*apiAT5-3* parasites were grown continuously in DMEM supplemented with 2.5 mM L-Tyr. TATi (38), TATi/Tomato, TATi/Δ*ku80* (39), and RHΔ*hxgprt*/Tomato (22) parasites were used as parental strains for the genetically modified parasites generated in this study.

### Generation of genetically modified *T. gondii* parasites

Guide RNA (gRNA)-encoding sequences specific to target genes were introduced into the vector pSAG1::Cas9-U6::sgUPRT (Addgene plasmid # 54467; (18)) using Q5 site-directed mutagenesis (New England Biolabs) as described previously (18). A list of the forward primers used to generate gRNA-expressing vectors for introducing frame-shift mutations are described in Table S4. In each instance, the reverse primer 5’-AACTTGACATCCCCATTTAC was used. For generating ‘knockout’ frameshift mutations in *Tg*ApiAT genes, gRNAs were designed to target the open reading frames of *Tg*ApiAT genes, and transfected into parasites on a vector that also expressed Cas9-GFP. Transfections were performed as described previously (40). GFP positive parasites were selected and cloned using flow cytometry 2-3 days following transfection using a FACSAria I or FACSAria II cell sorter (BD Biosciences). The region of the candidate genes targeted by the gRNAs were sequenced in clonal parasites, and clones in which the target gene had been disrupted by a frameshift mutation or insertion of a premature stop codon (i.e. where the open reading frame was disrupted) were selected for subsequent analyses. For 3’ replacements, gRNAs were selected to target a region near the stop codon of the gene of interest, using the primers listed in Table S5. In addition, a donor DNA sequence encoding a 3x HA tag was amplified by PCR to contain 50 bp of flanking sequences homologous to the target gene either side of the stop codon. Template DNA encoding the 3x HA tag was generated as a gBlock (Integrated DNA Technologies), with the sequence listed in Table S6. Forward and reverse primers used to amplify the HA tag for each target gene are also listed in Table S6. gRNA-expressing vectors, which simultaneously encode Cas9 fused to GFP, were cotransfected into *T. gondii* parasites with the donor DNA sequence. 2-3 days after transfection, GFP-Cas9-expressing parasites were selected and cloned into wells of a 96-well plate by flow cytometry as described above.

3’ replacement plasmids were created to epitope tag *Tg*ApiAT2, *Tg*ApiAT3-2, *Tg*ApiAT3-3, *Tg*ApiAT5-3, *Tg*ApiAT6-1, *Tg*ApiAT6-2 and *Tg*ApiAT7-1 using conventional crossover recombination methods as described previously (41). Regions of DNA homologous to the 3’ ends of the genes were amplified by PCR using primers described in Table S7, and ligated into the *Bgl*II and *Avr*II sites of the vector pgCH (5) or the *PacI* and *AvrII* sites of pLIC-HA_3_-DHFR (41). Resulting plasmids were linearized in the flanking sequence using restriction enzymes (Table S7), then transfected into TATi/Δ*ku80* parasites. Parasites were selected on chloramphenicol or pyrimethamine as described (40). In cases where we were unable to subsequently detect protein of approximately the expected molecular mass by western blotting, we confirmed correct integration of the HA tag by sequencing of CRISPR-modified 3’ ends (*Tg*ApiATs 5-1, 5-2, 5-4, 5-5 and 5-6; not shown), or by PCR screening (*Tg*ApiAT7-1). For assessing HA integration into the *Tg*ApiAT7-1 locus, DNA was extracted from *Tg*ApiAT7-1-HA parasite clones, and used as template in a PCR with the primers 5’-GGCGAAGAGAAGGCGTTG and 5’-GTCATCCCTTTTCTTCGATAA, with the presence of a 2.5 kb band indicative of successful integration (Fig S3C).

To complement the Δ*apiAT2* mutant with a constitutively-expressed copy of *Tg*ApiAT2, we amplified the open reading frame of *Tg*ApiAT2 from genomic DNA with the primers 5’-GATCGGATCCAAAATGGCGGCTGCTCAG and 5’-GATCCCTAGGCACAGCGACCTCTGGACTCGGT. We digested the resultant PCR product with *Bam*HI and *Avr*II and ligated this into the *Bgl*II and *Avr*II sites of the pUgCTH_3_ vector (5). The resultant vector was linearised with *Mfe*I, transfected into Δ*apiAT2* parasites and selected on chloramphenicol. To complement the Δ*apiAT5-3* mutant with a constitutively-expressed copy of *Tg*ApiAT5-3, we amplified the open reading frame of *Tg*ApiAT5-3 with the primers 5’-GATCGGATCCAAAATGGAGTCGACCGAGGCGACTAT and 5’-GATCCCTAGGCAGCACCTTCGGGACTTTTCTCTTC, using the *Tg*ApiAT5-3-expressing oocyte vector (described below) as template. We digested the resultant PCR product with *Bam*HI and *Avr*II and ligated this into the *Bgl*II and *Avr*II sites of the pUgCTH_3_ vector. The resultant vector was linearised with *Mfe*I, transfected into Δ*apiAT5-3* parasites and selected on chloramphenicol.

### Immunofluorescence assays and western blotting

Immunofluorescence assays and western blotting were performed as described previously (5). For western blotting, membranes were probed with rat anti-HA antibodies (clone 3F10, Sigma) at dilutions between 1:1,000 to 1:3,000, mouse anti-GRA8 (a kind gift from Gary Ward, U. Vermont, (42)) at a dilution of 1:80,000, or rabbit anti-*Tg*Tom40 antibodies (43) at 1:2,000 dilution, and horseradish peroxidase (HRP)-conjugated goat anti-rat (sc-2006, Santa Cruz Biotechnology), HRP-conjugated goat anti-mouse (sc-2005, Santa Cruz Biotechnology), or HRP-conjugated goat anti-rabbit (sc-2004, Santa Cruz Biotechnology) antibodies at dilutions of 1:5,000 to 1:10,000.

For immunofluorescence assays, samples were probed with the following primary antibodies: rat anti-HA (clone 3F10, Sigma) at a 1:200 dilution, mouse anti-P30 (clone TP3, Abcam) at a 1:2,000 dilution, rabbit anti-P30 (a kind gift from John Boothroyd, Stanford U) at dilutions between 1:25,000 and 1:90,000, or rabbit anti-GFP (a kind gift from Alex Maier, ANU) at a 1:200 dilution. Samples were next probed with the following secondary antibodies: CF488A-conjugated goat anti-rat (SAB4600046, Sigma) at a dilution of 1:500, AlexaFluor 488-conjugated goat anti-rat (4416, Cell Signaling Technology) at dilution of 1:250, AlexFluor488-conjugated goat anti-rabbit (A11008, Life Technologies) at a dilution of 1:500, AlexFluor546-conjugated goat anti-rabbit (A11035, Life Technologies) at a dilution of 1:500, AlexFluor546-conjugated goat anti-mouse (A11030, Life Technologies) at a dilution of 1:500, or AlexFluor647-conjugated goat anti-mouse (A21236, Life Technologies) at a dilution of 1:500.

Fluorescence microscopy was performed on a DeltaVision Elite system (GE Healthcare) using an Olympus IX71 inverted microscope with a 100X UPlanSApo objective lens (NA 1.40). Images were recorded using a CoolSNAP HQ2 camera. Images were deconvolved using SoftWoRx Suite 2.0 software, and images were linearly adjusted for contrast and brightness.

### Parasite growth and virulence assays

To measure parasite growth by plaque assays, either 150 parasites were added to wells of a 6-well plate containing confluent HFFs, or 500-1,000 parasites were added to confluent HFFs in 25 cm^2^ tissue culture flasks. Parasites were allowed to grow for 8 to 18 days before fixation and staining with crystal violet as described previously (40).

Fluorescence growth assays were performed as described previously (44, 45), with slight modifications. Briefly, wells of an optical bottom 96-well plate containing confluent HFFs were washed twice in medium lacking L-Tyr, L-Phe or L-Trp. Wells were filled with medium containing a range of L-Tyr, L-Phe or L-Trp concentrations. 2,000 parasites were inoculated into each of these wells, and plates were incubated at 37°C in a 5% CO_2_ incubator. Well fluorescence was measured 5 days post-inoculation in a FluoStar Optima fluorescence plate reader (BMG Labtech), a time point at which WT parasites were in mid-logarithmic stage of growth. Relative growth was expressed as a percentage of the well fluorescence in the optimum amino acid concentration for WT or Δ*apiAT5-3* parasites at this time point.

To measure parasite virulence, freshly egressed WT (RHΔ*hxgprt*/Tomato) or Δ*apiAT5-3* in RH/Δ*hxgprt*/Tomato parasites were filtered through a 3 μm polycarbonate filter, washed once in phosphate-buffered saline (PBS), then were diluted to 10^4^ parasites/ml in PBS. 10^3^ parasites were injected intraperitoneally into 7-week-old, female Balb/c mice using a 26-gauge needle. Mice were weighed regularly, and monitored for symptoms of toxoplasmosis (weight loss, ruffled fur, lethargy and hunched posture). Mice exhibiting terminal symptoms of toxoplasmosis were euthanized in accordance with protocols approved by the Australian National University Animal Experimentation Ethics Committee (protocol number A2016/42).

### [^13^C]Amino acid labelling and detection

Freshly egressed WT or Δ*apiAT5-3* tachyzoites (10^8^) were incubated in 500 μL of amino acid-free Roswell Park Memorial Institute 1640 medium supplemented with 2 mg/ml algal [^13^C]amino acid mix (Cambridge Isotope Laboratories) for 15 minutes at 37°C in a 5% CO_2_ incubator. [^13^C]amino acid labelling was terminated by rapid dilution in 14 ml of ice cold PBS. Parasite metabolites were extracted in chloroform:methanol:water (1:3:1 v/v/v) containing 1 nmol *sycllo*-inositol (Sigma). The aqueous phase metabolites were dried in a heated speedvac concentrator, methoxymated by treatment with 20 mg/ml methoxyamine in pyridine overnight, then trimethylsilylated by treatment with N,O-bis(trimethylsilyl)trifluoroacetamide containing 1% trimethylsilyl for 1 hr at room temperature. Samples were analyzed using GC-MS as described previously (46). The fractional labelling of all detected amino acids was estimated as the fraction of the metabolite pool containing one or more ^13^C-atoms after correction for natural abundance. Total metabolite counts were normalized to *scyllo*-inositol as an internal standard.

### *Xenopus laevis* oocyte preparation and *Tg*ApiAT5-3 expression

The open reading frame of *Tg*ApiAT5-3 was amplified from RHΔ*hxgprt* strain cDNA template using the primers 5’-GATCACCGGTCCACCATGGAGTCGACCGAGGCGACTAT and 5’-GATCCCTAGGCAGCACCTTCGGGACTTTTCTCTTC. The resultant product was digested with *Age*I and *Avr*II, and ligated into the *Xma*I and *Avr*II sites of the vector pGHJ-HA (5). The plasmid was linearised by incubation in *Not*I overnight, and complementary RNA (cRNA) encoding HA-tagged *Tg*ApiAT5-3 was prepared for injection into oocytes as previously described (47–49). *Xenopus laevis* oocytes were surgically removed and prepared for cRNA injection as described (48). For all transporter assays in oocytes, 15 ng of *Tg*ApiAT5-3 cRNA was micro-injected into stage 5 or 6 oocytes using a Micro4^TM^ microsyringe pump controller and A203XVY nanoliter injector (World Precision Instruments). Maintenance of animals and preparation of oocytes was approved by the Australian National University Animal Experimentation Ethics Committee (protocol number A2014/20).

### Oocyte surface biotinylation and whole membrane preparation

Oocyte surface biotinylation and whole membrane preparations were performed as described previously (48, 50). Briefly, for surface biotinylation, 15 oocytes were selected 3-6 days post cRNA injection, washed thrice in ice-cold PBS (pH 8.0), incubated for 45 mins at room temperature in 0.5 mg/ml of EZ-Link^TM^ Sulfo-NHS-LC-Biotin (Thermo Fisher Scientific), and then washed thrice more in ice-cold PBS. Oocytes were subsequently solubilised in oocyte lysis buffer (20□mM Tris-HCl pH 7.6, 150□mM NaCl, 1% v/v Triton X-100) for 2 hr on ice. Samples were centrifuged at 16,000 *g*, and the supernatant was mixed with 50 μl of streptavidin-coated agarose beads (Thermo Fisher Scientific). The mixture was incubated at 4°C on slow rotation overnight. Beads were washed 4 times with oocyte lysis buffer before elution in SDS-PAGE sample buffer. For whole membrane preparation, 10-25 oocytes were homogenised by trituration in homogenisation buffer (50 □mM Tris-HCl pH 7.4, 100 □mM NaCl, 1 mM EDTA, protease inhibitors). Homogenised oocytes were centrifuged at 2,000 *g* for 10 min at 4 °C and the resulting supernatant further centrifuged for 30 min at 140,000 *g* at 4°C. The resulting pellet was washed with homogenisation buffer and solubilised in homogenisation buffer containing 4 % (w/v) SDS, and then in SDS-PAGE sample buffer. Protein samples from surface biotinylation and whole membrane preparations were separated by SDS-PAGE then detected by western blotting as described above.

### Oocyte uptake, efflux and electrophysiology

For uptake experiments in either non-preloaded, preloaded, or pre-injected oocytes, batches of 10 oocytes were washed 4 times in ND96 buffer (96 mM NaCl, 2 mM KCl, 1 mM MgCl2, 1.8 mM CaCl_2_, 5 mM HEPES, pH 7.4) at RT, and then incubated in the desired concentration of radiolabelled substrates as indicated in figure legends. For all substrate screening measurements, uptake was measured over 10 mins. For kinetic experiments, parallel batches of oocytes were preloaded with 0 – 1.78 mM L-Tyr, and uptake of [^14^C]Tyr at a range of L-Tyr concentrations was measured over 10 mins. For other uptake experiments, uptake was measured for the time-course indicated in the figures. Uptake was quenched by washing oocyte batches four times in ice-cold ND96.

For efflux experiments, batches of 5 oocytes/substrate were preloaded with [^14^C]L-amino acids as described below. Following preloading, oocytes were washed 4 times in ND96 and incubated in trans-stimulating substrates at concentrations described in the figure legends. To measure the amount of efflux, oocytes were incubated in 500 μl aliquots of the extracellular solution, of which 100 μl was removed for each time point and the efflux immediately quenched by washing oocytes in four times in ice-cold ND96. Oocyte retention was measured by removing the oocytes following quenching of efflux. For the substrate efflux screen depicted in Fig 6B, the extracellular solution was sampled at 5 min to ensure initial rate measurements. For efflux experiments depicted in Fig 5C, efflux was measured for the time-course indicated in the figure.

Following all uptake and efflux experiments, oocytes or aliquots were distributed into OptiPlate96-well plates (Perkin-Elmer) and oocytes were lysed overnight in 10 % (w/v) SDS. 150 μl/well of Microscint-40 scintillation fluid (Perkin-Elmer) was added to the samples, and plates covered and shaken for 5 min before radioactivity was counted on a Perkin-Elmer MicroBeta^2^ 2450 microplate scintillation counter.

All steady-state electrical recordings were made with an Axon GeneClamp 500B amplifier (Axon Instruments) in a two-voltage clamp configuration as previously described (50, 51). Voltage clamp was set to –50 mV or 0 mV and data were sampled at 3 Hz using pClamp 8.2 software (Axon Instruments). Boron silicate microelectrodes capillaries (World Precision Instruments) with a tip resistance of: 1.5 ≥ *R_e_* ≥ 0.5 MΩ were pulled by a P-97 Flaming/Brown micropipette puller (Sutter Instruments) and filled with 3 M KCl. Silver microelectrodes were coated using a 5 M NaCl single-chamber galvanic cell to form AgCl_2_ electrodes. The membrane potential was adjusted digitally in voltage-clamp mode between 0 and 50 mV. Oocytes were chosen for recording when they had a resting membrane potential –25 mV < E_m_ < –45 mV. ND96 (pH 7.4) was used as the control solution for all electrophysiological recordings. Assay buffer pH was varied by mixing different ratios of acidic ND96 (pH 3.6) (5 mM MES instead of HEPES) with basic ND96 (pH 10) (5 mM Tris instead of HEPES).

### Oocyte preloading and pre-injection of trans-substrates

For uptake experiments measuring *trans*-stimulation by a range of amino acids and other metabolites (Fig 6A and Fig S5B), all L-amino acids substrates and metabolite mixes, except for L-Tyr, were pre-injected at 25 nl/oocyte using a Micro4^TM^ micro-syringe pump controller and A203XVY nanoliter injector (World Precision Instruments). All pre-injected oocytes were incubated on ice for 30 mins prior to uptake experiments. Stock solutions containing 100 mM L-amino acids in ND96 were pre-injected to give a calculated cytosolic concentration of 5 mM, based on an assumed free aqueous volume of 500 nl/oocyte. Stage 5 or 6 oocytes diameters vary significantly from 1 – 1.3 mm and free aqueous oocyte volumes measured from 368 to > 500 nl (52, 53). Therefore, calculations of cytosolic concentrations from pre-injection should be treated as approximations only. In Fig. S5B, pre-injection of different metabolite groups was conducted to give estimated final concentrations of each metabolite as indicated in Table S2. The low solubility of L-Tyr in aqueous solutions (0.453 g/L at 25°C, pH 7.4; (54)) necessitated preloading, rather than pre-injecting, substrate for *trans*-stimulation and efflux substrate specificity experiments. L-Tyr solutions were made by dissolving 2.5 mM L-Tyr in ND96 at 37°C and performing dilutions in ND96 to the required concentrations. Pre-loading of L-tyrosine in *Tg*ApiAT5-3-injected oocytes was tested by timing the pre-loading of 2.5 or 1 mM [^14^C]Tyr (data not shown). L-tyrosine equilibrium was reached after 10 to 12 hr, while uninjected oocytes reached a similar cytosolic concentration after incubation of approximately 68-72 hr. As a consequence, L-Tyr was preloaded for 32 hr in *Tg*ApiAT5-3 injected oocytes and for 72 hr in uninjected oocytes for all *trans*-stimulation experiments. Greater than 90% of [^14^C]Tyr was effluxed from oocytes preloaded with 2.5 mM labelled L-Tyr, indicating L-Tyr is not significantly metabolised during the preloading times used in experiments (data not shown).

All efflux substrate screening with [^14^C]labelled amino acids (Fig 6B) were conducted by preloading [^14^C]labelled L-amino acids for 3 hr prior to uptake. [^14^C]Tyr was preloaded at a concentration of 2.5 mM, while the other [^14^C]labelled L-amino acids were preloaded at a concentration of 5 mM. Calculation of the pre-loaded [^14^C]labelled L-amino acids concentrations were conducted using a control set of oocytes for each substrate, pre-loaded in parallel to those used for efflux *trans*-stimulation. Calculations were made assuming a cytosolic volume of 500 nl/oocyte.

### Oocyte data analysis and statistics

All oocyte data were analyzed using OriginPro (2015). All data displayed in figures represent the mean ± S.D. except where otherwise indicated. Unless uptake data from uninjected oocytes is included in figures, uptake in uninjected oocytes was subtracted from uptake in *Tg*ApiAT5-3-injected oocytes to give the ‘ *Tg*ApiAT5-3-mediated uptake’. All data sets were analysed for Gaussian normalcy by first running a Shapiro-Wilk test prior to analysis and used only if passing the normalcy test at the P < 0.05 level. Multi-variant experiments with 3 or more experimental conditions were subjected to a one-way ANOVA with Dunnet’s post-hoc test and significance tested at the P < 0.05 level.

Time-course analysis of uptake and oocyte retention data of L-Tyr in *Tg*ApiAT5-3-injected oocytes were fitter to 1^st^ order integrated rate equations:

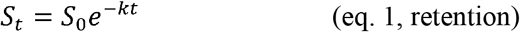

Or:

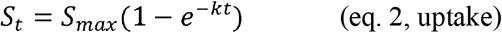

Equation 2 being the Box Lucas 1 model with zero offset (55), where S_t_, and S_0_, and are the amount of substrate (S) at variable time (t), or when t = 0, S_max_ is the vertical asymptote of substrate amount, and k is the 1^st^ order rate constant.

Steady-state kinetic data collected under initial rate conditions were fitted to both the Michaelis-Menten equation:

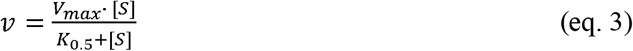

And a Scatchard linear regression equation:

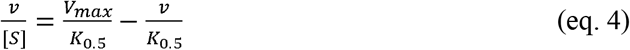

In the Scatchard regression, the apparent Michaelis constant (K_0.5_) is derived from the slope (1/–K_0.5_) and the maximal rate (V_max_) from the ordinate intercept (V_max_/K_0.5_).

All curve fittings were evaluated using adjusted R^2^ values as indicated in the text and figure legends. All non-linear fitting was conducted using the Levenburg-Marquardt algorithm, with iteration numbers varying from 4 to 11 before convergence was attained.

### *T. gondii* amino acid assays

Amino acid uptake assays were carried out as described previously (5), with slight modifications. Briefly, extracellular parasites were washed twice in Dulbecco’s PBS pH 7.4 (Sigma) supplemented with 10 mM glucose (PBS-glucose). Parasites were incubated in PBS-glucose containing radiolabelled amino acids, and 200 μl aliquots removed at regular time points. Parasite samples were centrifuged through an oil mix to separate parasites from unincorporated radiolabel as described previously (5). L-Arg uptake was measured by incubation in 0.1 μCi/mL [_^14^_C]Arg and 100 μM L-Arg, L-Tyr uptake was measured by incubation in 0.25 μCi/mL [_^14^_C]Tyr and 60 μM L-Tyr, and L-Phe uptake was measured by incubation in 0.1 μCi/mL [_^14^_C]Phe and 15 *μ*M L-Phe. The time courses of radiolabel uptake in each amino acid tested were fitted by a single exponential function and the initial rate of transport was estimated from the initial slope of the fitted line.

## Acknowledgements

We are grateful to students from the 2016 Biology of Parasitism course (Woods Hole, MA) for performing some of the early studies to characterise the role of *Tg*ApiAT5-3 in aromatic amino acid transport. We thank Harpreet Vohra and Michael Devoy for performing flow cytometry.

---

**Fig S1. Multiple sequence alignment of ApiAT family proteins from apicomplexans and chromerids**. A multiple sequence alignment of the 67 ApiAT family proteins examined in this study. The alignment is presented as a “fingerprint”, where each residue is represented by a thin vertical line that has been shaded to represent the degree of conservation (as described previously; (58)). Residues with >70 % identity in the ApiAT alignment are depicted in purple, residues with 50-70 % identity are depicted in cyan, residues where > 50% of residues have similar amino acids or where amino acids are similar to residues in the above identity groupings are depicted in magenta, non-conserved residues are depicted in grey, and gaps in the sequences are white. The approximate locations of the predicted transmembrane domains are represented by numbered bars, and the location of the MFS signature sequence between transmembrane domains two and three has been highlighted.

**Fig S2. Multiple sequence alignment of a selection of ApiAT family proteins from apicomplexans with human LAT3 and LAT4 proteins.** A multiple sequence alignment of ApiAT-family proteins from apicomplexans (*Tg*ApiAT1, *Tg*ApiAT2, *Pb*ApiAT8-1, *Tg*ApiAT6-1 and *Tg*ApiAT5-3) and the human LAT3 and LAT4 proteins (*Hs*LAT3 and *Hs*LAT4). Residues with >70% sequence identity are shaded in black and residues with >70% sequence similarity are shaded in gray. The red box highlights the MFS signature sequence.

**Fig S3. Genetic modifications to introduce HA tags into the native loci of *Tg*ApiAT genes in *T. gondii*.** (A) Single cross-over recombination approach, where a vector containing a homologous flanking sequence to the target gene, in addition to a chloramphenicol resistance marker (ChlR), is introduced into *T. gondii* parasites. Single cross-over recombination results in the insertion of a HA tag into the 3’ region of the open reading frame of the target gene. The approximate position of the primers used to screen *Tg*ApiAT7-1-HA clones are depicted. (B) CRISPR/Cas9 genome editing approach, where a guide RNA (gRNA) is designed to target a region near the stop codon of the target gene. When co-expressed with Cas9-GFP, the gRNA mediates a double-stranded break in the parasite genome near the stop codon of the target gene. The gRNA/Cas9-GFP vector is co-transfected with a donor DNA product that contains a HA tag flanked on either side with 50 bp of sequence homologous to regions immediately up and downstream of the stop codon in the target gene. Homologous repair results in introduction of the HA tag into the 3’ region of the open reading frame of the target gene. (C) PCR screen to test for integration of the HA tag into the *Tg*ApiAT7-1 locus. The presence of a 2.5 kb band that is absent from the wild type (WT) control indicates that clones 2-5 have successfully integrated the HA tag.

**Fig S4. Characterisation of *Tg*ApiAT5-3 expressed in oocytes** (A) Western blot with anti-HA antibodies on whole membrane preparations (WMP) and surface biotinylated proteins (SB) in oocytes expressing HA-tagged *Tg*ApiAT5-3 (5–3) or oocytes that were uninjected (u.i.). (B) Efflux and retention of preloaded [^14^C]Tyr in uninjected oocytes. Uninjected oocytes were preloaded by incubation in 1 mM [^14^C]Tyr for 72 hr as described in methods. Subsequent efflux (filled shapes) and retention (open shapes) of the preloaded labelled substrate was measured over the time-course indicated in the presence in the extracellular buffer of 2.5 mM L-Tyr (squares) or in the absence of L-Tyr (circles). Data show the mean efflux and retention in 5 oocytes from a single experiment ± standard deviation, and are representative of 3 independent experiments. (C) *Tg*ApiAT5-3-expressing oocytes (black) or uninjected oocytes (white) were preloaded via incubation in 2.5 mM L-tyrosine for 32 or 72 hr, respectively, as described in methods. Subsequent uptake of 1 mM [^14^C]Tyr was measured in buffer where the ions were replaced as indicated. For Na+ replacement conditions, the replacement cation is written at the top of the respective histogram. Data show the mean uptake in 10 oocytes from a single experiment ± standard deviation, and are representative of 3 independent experiments. Uptake in *Tg*ApiAT5-3-expressing oocytes was not significantly different in any condition tested (P > 0.05, one-way ANOVA, Dunnet’s post-hoc test). (D) *Tg*ApiAT5-3-expressing oocytes were impaled and recorded using a two-voltage clamp amplifier configuration 4-5 days post-cRNA injection. Oocytes were continuously perfused with gravity-fed ND96 buffer (pH 7.4) until otherwise indicated by the arrows in the current tracings. Top: representative current trace upon the addition of 1 mM L-Tyr at E_m_ = –50 mV or 0 mV. Bottom: representative current trace upon the change to pH 9.0 and incubation in 1 mM L-Tyr. No baselines were corrected in either tracing. Data are representative of 12 replicates.

**Fig S5. Substrate specificity of *Tg*ApiAT5-3** (A) Uptake of 500 μM [^14^C]Tyr was measured in *Tg*ApiAT5-3-expressing oocytes (black) or uninjected oocytes (white) over 10 mins in presence of 500 μM unlabelled L-amino acids. Data show the mean uptake in 10 oocytes from a single experiment ± standard deviation, and are representative of 2 independent experiments (*, P > 0.05, one-way ANOVA, Dunnet’s post-hoc test. Where significance values are not shown, the differences are not significant, P > 0.05). (B) Uptake of [^14^C]Tyr in *Tg*ApiAT5-3-expressing oocytes (black) or uninjected oocytes (white) over 10 mins, where oocytes were pre-injected with uptake buffer (ND96), preloaded with 2.5 mM L-Tyr, or pre-injected with various substrate mixes, including L-amino acids (L-AA1-3), amino acid derivatives (AA derivatives 1-3), D-amino acids (D-AA), nucleosides, nitrogen bases, or sugars (see Table S2 for compositions). Data show the mean uptake in 8-10 oocytes from a single experiment ± standard deviation, and are representative of 3 independent experiments (*, P < 0.05, one-way ANOVA, Dunnet’s post-hoc test. Where significance values are not shown, the differences are not significant, P > 0.05). (C) Uptake of various [^14^C]Amino acids (at 1 mM concentration) was measured in *Tg*ApiAT5-3-expressing oocytes (black) or uninjected oocytes (white) over 10 mins. Data show the mean uptake in 10 oocytes from a single experiment ± standard deviation, and are representative of 3 independent experiments (*, P > 0.05, one-way ANOVA, Dunnet’s post-hoc test, for differences between *Tg*ApiAT5-3-injected and uninjected oocytes for the same substrate. Where significance values are not shown, the differences are not significant, P > 0.05).

**Fig S6. Time courses for the uptake of [^14^C]Tyr, [^14^C]Phe and [^14^C]Arg in *T. gondii*.** Uptake of [^14^C]Tyr (A), [^14^C]Phe (B), and [^14^C]Arg (C) in WT (A-C), Δ*apiAT5-3* (A-C), and c*Tg*ApiAT5-3/Δ*apiAT5-3* (A, C) strain parasites. Uptake was measured in PBS-glucose containing either 60 μM unlabelled L-Tyr and 0.1 μCi/mL [^14^C]Tyr (A), 30 μM unlabelled L-Phe and 0.1 μCi/mL [^14^C]Phe (B), or 100 μM unlabelled L-Arg and 0.1 μCi/mL [^14^C]Arg (C). Data points represent the mean ± SEM from three independent experiments. Lines represent fitted single-order exponential curves, from which the initial rates were calculated and depicted in Fig 7.

**Fig S7. Complementation of Δ*apiAT5-3* strain parasites with a constitutive copy of *Tg*ApiAT5-3 restores parasite growth in DMEM.** Plaque assays depicting growth of TATi/Tomato (WT) parasites (top), Δ*apiAT5-3* parasites (middle), and Δ*apiAT5-3* parasites complemented with a constitutive copy of *Tg*ApiAT5-3 (c*Tg*ApiAT5-3/Δ*apiAT5-3*; bottom). 500 parasites were added to 25 cm2 tissue culture flasks and cultured in DMEM (left) or DMEM containing 2.5 mM L-Tyr (right) for 11 days before fixation and staining with crystal violet. Data are representative of two independent experiments.

**Fig S8. *T. gondii* parasites are auxotrophic for all three proteinogenic aromatic amino acids.** Fluorescence growth assays measuring the growth of WT (black) and Δ*apiAT5-3* (gray) parasites in DMEM containing different concentrations of L-Tyr (A), L-Phe (B) and L-Trp (C). The growth of parasites is expressed as a percentage of the optimal concentration for each amino acid tested in each parasite strain, and was measured at mid-log phase for this optimal concentration (5 days post-inoculation). Parasite growth was determined using the same amino acid concentrations used in Fig 8, but included a 0 mM concentration (which was not possible to depict in Fig 8 because of the log scale on the x axis). For simplicity, only the following amino acid concentrations are depicted in this figure: 0 mM, 0.423 mM and 2.5 mM L-Tyr (A), 0 mM, 0.32 mM and 10 mM L-Phe (B), and 0 mM, 0.063 mM and 1 mM L-Trp (C). The data for 0.423 mM L-Tyr (the normal DMEM concentration of L-Tyr) were interpolated from curve fitting while 0.32 mM L-Phe and 0.063 mM L-Trp (the nearest tested concentrations to those present in DMEM) were experimental data points. Data represent the mean ± SEM from three independent experiments.

**Fig S9. Preliminary characterisation of the Δ*apiAT5-3*/RHΔ*hxgprt*/Tomato mutant** (A) Uptake of [^14^C]Tyr in RHΔ*hxgprt*/Tomato (WT, blue) and Δ*apiAT5-3* in RHΔ*hxgprt*/Tomato (red) parasites. Uptake was measured in PBS-glucose containing 60 μM unlabelled L-Tyr and 0.1 μCi/mL [^14^C]Tyr. Data points represent the mean from a single replicate. Lines represent fitted single-order exponential curves. (B) Fluorescence growth assay for Δ*apiAT5-3*/RHΔ*hxgprt*/Tomato parasites cultured for 16 days (when parasites were in mid-logarithmic stage) in DMEM containing a range of L-Tyr concentrations. Growth is expressed as a percentage of maximum growth measured on day 16, and data points represent the mean from three technical replicates (± standard deviation) in a single experiment. The concentration of L-Tyr in standard DMEM is depicted by a dashed line. (C) Plaque assays depicting growth of RHΔ*hxgprt*/Tomato (WT) parasites and Δ*apiAT5-3*/RHΔ*hxgprt*/Tomato parasites in normal DMEM (top) or DMEM containing 2.5 mM L-Tyr. 1000 parasites were added to each 25 cm^2^ tissue culture flask and incubated for 8 days (left and centre) or 18 days (right) before developing. Data are from a single independent experiment.

